# Combining an autoactivated cGAS mutant with membrane-tethered IL-12 via mRNA lipid nanoparticles promotes systemic tumor control

**DOI:** 10.64898/2026.10.15.759215

**Authors:** Anli Zhang, Yan Fang, Ruoshi Peng, Minglu Yan, Zihang Chen, Tuo Li, Jiaxing Tan, Lijun Sun, Zhijian J. Chen

**Author notes:** These authors contributed equally (Anli Zhang, Yan Fang).

## Abstract

Local tumor therapies rarely induce systemic immunity capable of controlling metastatic disease. Here, we engineer an mRNA lipid nanoparticle (LNP) platform to coordinate innate immune sensing and T cell activation in situ through co-delivery of an autoactivated cyclic GMP-AMP synthase mutant (cGAS^R241E^) and membrane-tethered IL-12 (IL-12TM). cGAS^R241E^ mRNA LNPs autonomously generated cyclic GMP-AMP *in vivo*, activating host STING signaling and initiating antitumor responses. Co-administration with IL-12TM mRNA LNPs promoted localized cytokine production, reduced regulatory T cells, and restored effector function in tumor-infiltrating T cells. This combination elicited systemic antitumor immunity and suppressed distal tumors across multiple mouse models. The therapeutic efficacy required CD8^+^ T cells and IFN-γ signaling in host, supporting a model in which innate DNA sensing is coupled to IL-12-mediated T cell functional reinforcement. In addition, the combination enhanced responsiveness to PD-L1 blockade. Together, these findings establish a clinically translatable strategy to integrate cGAS-STING activation with cytokine-driven T cell programming to drive systemic tumor control.

## Main

While cancer immunotherapy has revolutionized cancer treatment, only a subset of patients derives durable benefit, with the majority relapsing due to primary or acquired resistance. There is therefore an urgent need to develop novel immunotherapeutic strategies capable of eliciting robust and systemic antitumor immunity. Tumor derived double-stranded DNA can be sensed by cyclic GMP-AMP synthase (cGAS), which catalyzes the conversion of GTP and ATP into the second messenger 2′3′-cyclic GMP-AMP (cGAMP). cGAMP binds and activates the adaptor protein STING (Stimulator of Interferon Genes), leading to phosphorylation of TBK1 and IRF3 and subsequent induction of type I interferons (IFNs) and proinflammatory cytokines^1, 2, 3, 4, 5^. Activation of the cGAS-STING pathway enhances tumor antigen presentation by antigen-presenting cells, especially dendritic cells (DCs)^6, 7, 8^, promotes T cell activation and differentiation^9, 10^ and reprograms the immunosuppressive tumor microenvironment (TME)^11, 12^, which eventually enhance antitumor immunity in preclinical models. However, clinical evaluation of STING agonists has been limited by transient activity and systemic toxicity, constraining their therapeutic window^13, 14^.

The mRNA lipid nanoparticle (LNP) platform has unique advantage by enabling sustained protein expression while restricting cGAMP production within target cells, thereby limiting systemic exposure^15^. Recent studies have explored delivery of constitutively active STING and cGAS variants using mRNA-LNPs to enhance antitumor immunity. However, whether upstream DNA sensing can be strategically integrated with cytokine signaling to drive systemic tumor control remains unclear^16, 17, 18, 19^. In particular, activation of the cGAS-STING pathway alone is often insufficient to sustain robust effector T cell function within the TME, suggesting a need for synergistic signals that reinforce T cell immunity.

Interleukin-12 (IL-12) is a heterodimeric cytokine composed of p35 and p40 subunits and is produced primarily by antigen-presenting cells, including macrophages and DCs, under tight regulatory control^20, 21, 22, 23^. IL-12 promotes Th1 differentiation and enhances cytotoxic CD8^+^ T cells and natural killer (NK) cell activity through IFN-γ dependent mechanisms, showing potent antitumor activity in preclinical models^24, 25^. However, systemic IL-12 therapy in clinical trials has been associated with severe toxicities, including hematologic and hepatic adverse effects, limiting its therapeutic application^26, 27^. To mitigate these effects, strategies that confine IL-12 activity to the TME, including targeted delivery and membrane-tethered formats delivered by mRNA LNPs, have been developed^28, 29^. Although these approaches preserve local antitumor efficacy, they have generally failed to elicit robust abscopal responses, defined as suppression of untreated distal tumors.

Here, we began with *in vivo* cytokine screening and identified synergy between cGAMP-mediated innate immune activation and IL-12 signaling that drives systemic antitumor immunity. We then engineered LNPs encapsulating mRNAs encoding an autoactivated cGAS^R241E^ mutant together with membrane-tethered IL-12, enabling coordinated innate and cytokine signaling within the TME while minimizing systemic toxicity. This combination enhances control of both primary and distal tumors in two-tumor models and requires CD8^+^ T cells and IFN-γ signaling on host cells. Together, these findings establish an mRNA-LNP–based strategy to integrate innate DNA sensing and cytokine signaling to drive systemic tumor control.

## Results

### IL-12 synergizes with cGAMP to reduce both local and distal tumor burden

Because cGAS-STING signaling enhances innate immune sensing but does not directly promote adaptive immunity, and its indirect effects on T cell responses are often constrained by the TME, we reasoned that cytokines capable of activating or expanding effector T cells might synergize with cGAMP to achieve systemic antitumor efficacy. To test this, we used a B16F10 two-tumor model to evaluate both local and distal tumor control following combination treatment with cGAMP and cytokines known to directly modulate T cell function, including IL-2 (here we used a super-mutant IL-2 that preferentially targets CD8 T cells over regulatory T cells (Tregs)), IL-15/IL15Rα complexes, IL-21 and IL-12, all of which have demonstrated antitumor activity in preclinical or clinical settings^30, 31^.

Although IL-2 monotherapy produced moderate tumor growth inhibition, its combination with cGAMP did not significantly enhance therapeutic efficacy. (**Fig. 1a**). Similarly, neither IL-15/IL-15Rα nor IL-21 elicited significant antitumor effects as monotherapies, and their combination with cGAMP failed to improve local or distal tumor control (**Fig. 1b, c**). In contrast, combining IL-12 with cGAMP resulted in a marked enhancement of antitumor activity, with significant suppression of both treated (local) and untreated (distal) tumors (**Fig. 1d**), suggesting a functional synergy between STING activation and IL-12 signaling. These results identify IL-12 as the most effective cytokine partner among those tested for cGAMP-mediated innate immune activation. Notably, combination treatment was associated with reduced body weight gain, particularly in the cGAMP plus IL-12 group (**Extended Data Fig. 1**), suggesting a potential trade-off between efficacy and tolerability.

**Fig. 1.**
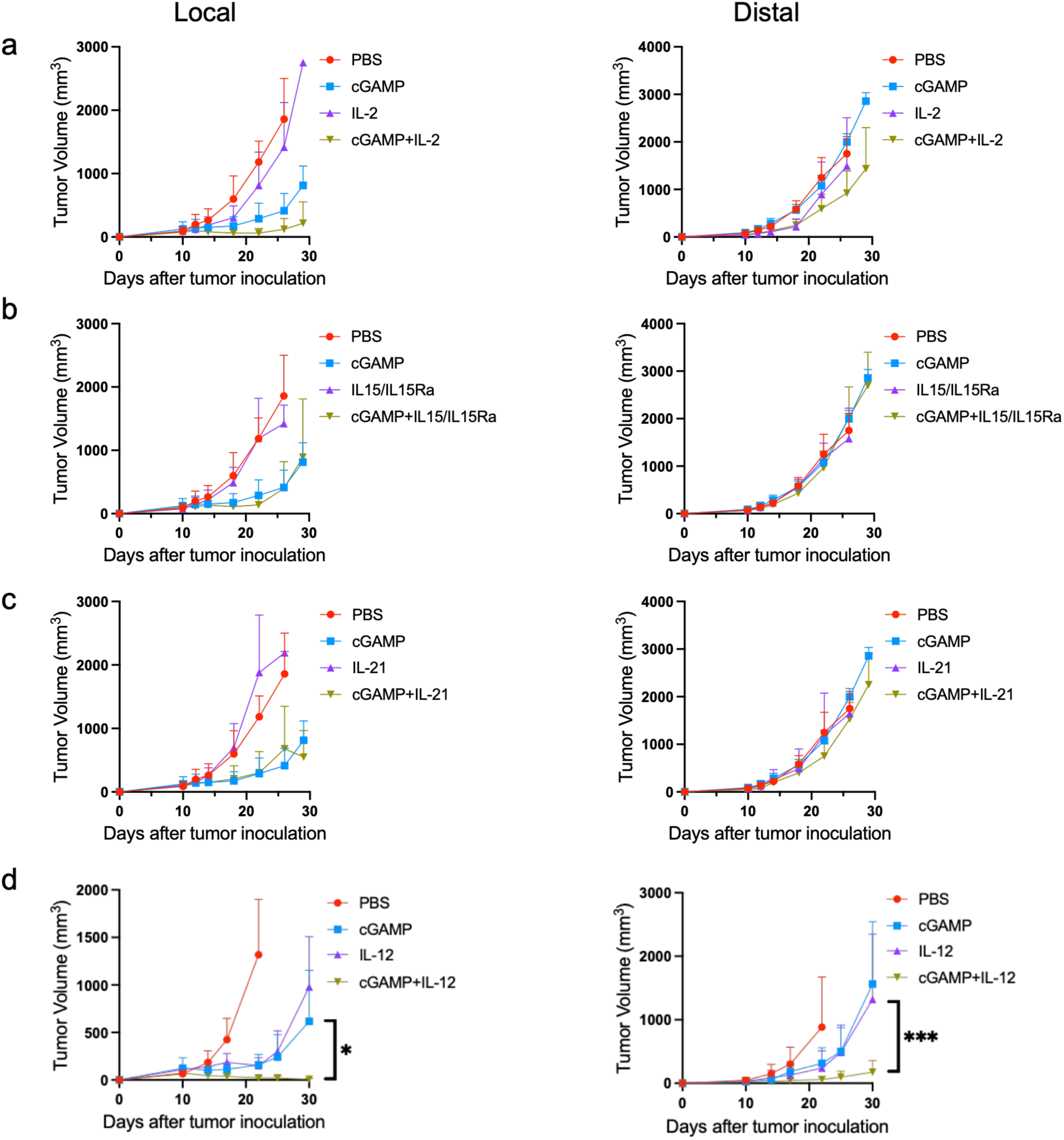
IL-12 synergizes with cGAMP to reduce both local and distal tumor burden. B16 tumor cells were inoculated into right dorsal flank (5×10^5^) and left dorsal flank (1×10^5^) on day 0. Mice were treated intratumorally on days 10, 14 and 18 with cGAMP (10 μg), IL-2 (500 ng) (a), IL-15 (80ng)/1L-15A (480 ng) (b), IL-21(500 ng) (c) or IL-12(30 ng) (d). Tumor growth was monitored. *p<0.05, ***p<0.001.

### Combination of cGAMP and IL-12 reprograms tumor-infiltrating T cells

To investigate how combined IL-12 and cGAMP treatment reshapes systemic T cell responses, we performed unbiased single-cell RNA sequencing (scRNA-seq) of tumor-infiltrating lymphocytes. Tumors from five control mice and eight cGAMP+IL-12-treated mice were pooled by treatment condition before library preparation. Because locally treated tumors regressed to very small sizes following combination therapy, insufficient cells were recovered for analysis; therefore, subsequent analysis focused on T cells infiltrating distal, untreated tumors. Unsupervised clustering identified ten distinct CD4⁺ T cell populations (**Extended Data Fig. 2a, c**). The pooled treated sample showed lower representation of the C6 Treg state and higher representation of the C8 activated Th1-like state relative to the pooled control sample (**Fig. 2a, b**). These findings indicate a potential shift from immunosuppressive CD4⁺ T cell states toward IFN-γ–producing effector phenotypes in distal tumors.

**Fig. 2.**
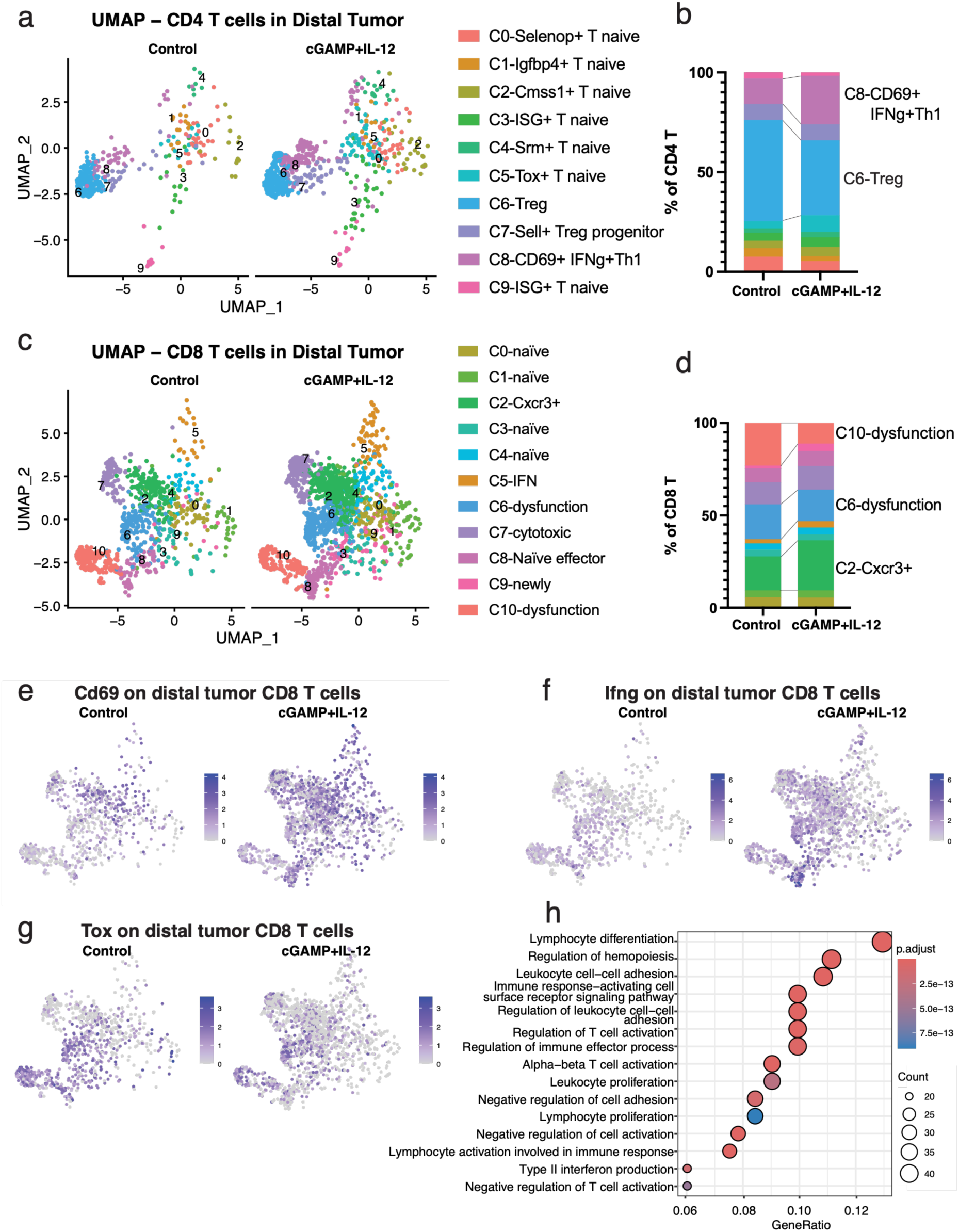
Combination of cGAMP and IL-12 reprograms tumor-infiltrating T cells. (a-d) UMAP plot shows unsupervised clustering and distribution of CD4^+^ (a) and CD8^+^ (c) T cells of distal tumors from control and cGAMP+IL-12 treated groups. Bar graph shows cell cluster distribution of CD4^+^ (b) and CD8^+^ (d) T cells from control and cGAMP+IL-12 treated tumors. (e-g) Cd69 (e), Ifng (f) and Tox (g) expression are highlighted in the CD8^+^ T cell UMAP plot. (h) GO enrichment analysis of CD8-C2 Cxcr3^+^ cells.

Eleven clusters were identified in CD8^+^ T cells (**Extended Data Fig. 2b, d**). The pooled treated sample showed increased representation of the C2 CXCR3⁺ effector-like state and reduced representation of two dysfunction-associated states, C6 and C10 (**Fig. 2c,** d). C2 was characterized by Cxcr3, Ly6c2, Cd69, and Ccl5, whereas C6 and C10 expressed dysfunction-associated genes including Tox, Pdcd1, Havcr2, Entpd1, Lag3, and Tigit. The major states remained identifiable across clustering resolutions and alternative embeddings, and the direction of the C2 increase and combined C6+C10 decrease was preserved across resolutions 0.4-1.4 (**Extended Data Fig. 3**). UMAP visualization further revealed increased activation markers *Cd69* and *Ifng* (**Fig. 2e, f**) and reduced *Tox* expression (**Fig. 2g**) in CD8^+^ T cells following combination treatment. Gene set enrichment analysis showed enrichment of T cell activation and type II IFN signaling pathways within the expanded C2 cluster (**Fig. 2h**), consistent with a functionally reinvigorated effector state. Collectively, these data indicate that IL-12 synergizes with cGAMP-mediated innate immune activation to reprogram tumor-infiltrating T cells toward activated, IFN-γ–competent effector phenotypes in distal tumors.

### cGAS^R241E^ mRNA LNPs induce STING-dependent type I IFN responses through sustained cGAMP production

Given the transient body weight loss observed following combined administration of cGAMP and IL-12 (**Extended Data Fig. 1k**), we sought to improve delivery efficiency while limiting systemic exposure by developing an mRNA LNP-based strategy. Under physiological conditions, cGAS activity is tightly regulated through chromatin tethering, which prevents aberrant activation by self-DNA. The autoactivated cGAS^R241E^ mutant disrupts this chromatin tethering, resulting in constitutive enzymatic activity and spontaneous cGAMP synthesis^32^. To evaluate its immunostimulatory potential, we generated mRNA constructs encoding either cGAS^R241E^ mutant or a catalytically inactive control (cGAS^E211A/D213A^, hereafter referred to as cGAS^Dead^ ^33^, and formulated them into LNPs (**Fig. 3a**). The resulting LNPs exhibited a hydrodynamic diameter of approximately 80 nm with a polydispersity index of ∼20%, indicating a homogeneous size distribution suitable for efficient cellular delivery (**Extended Data Fig. 4a, b**). Cryo-electron microscopy further confirmed the uniform particle formation (**Extended Data Fig. 4c**).

**Fig. 3.**
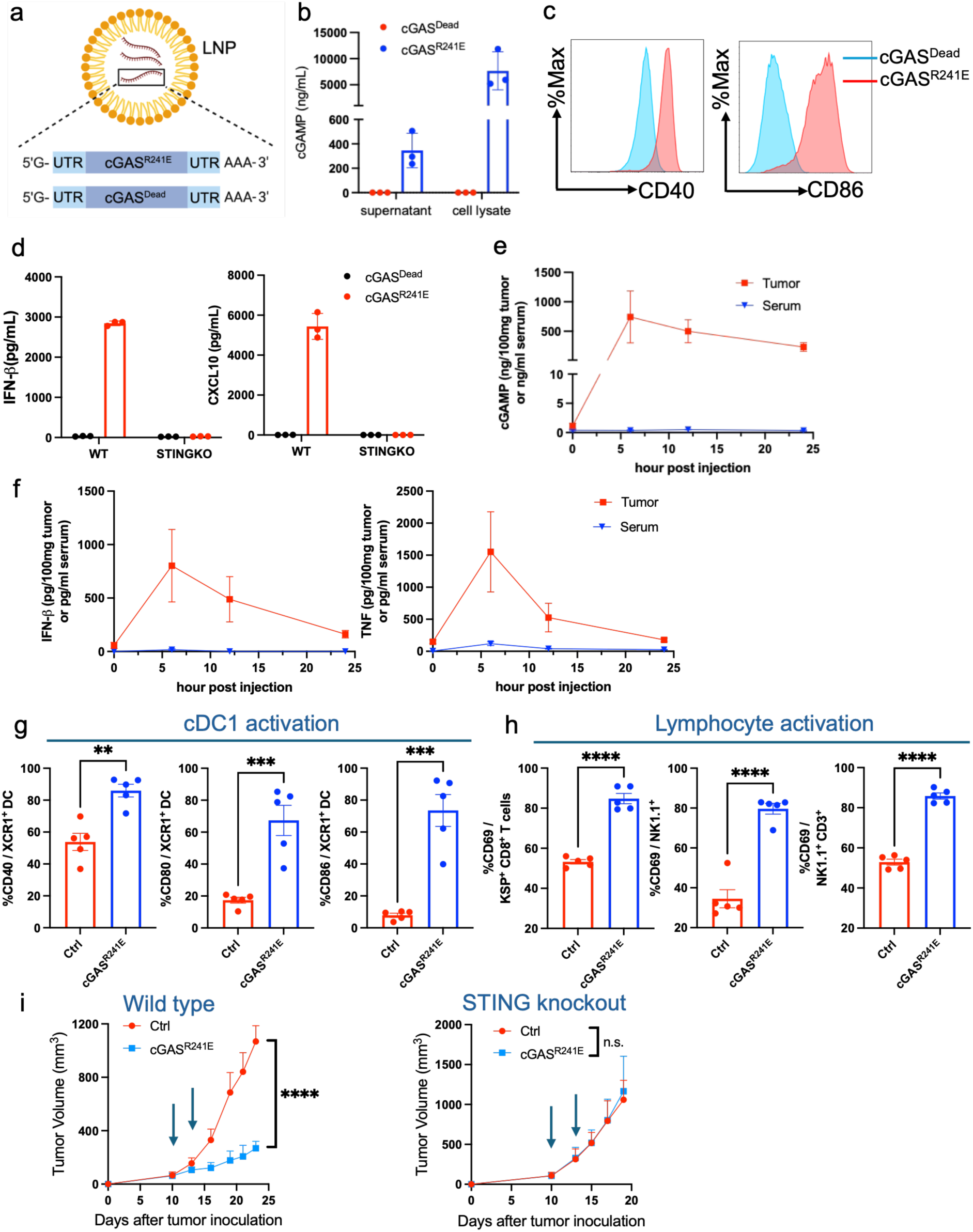
cGAS^R241E^ LNP spontaneously produces cGAMP and induces STING-dependent type I IFN production. (a) Schematic of the autoactivated cGAS^R241E^ mutant and catalytic dead cGAS^Dead^ mRNA constructs. (b) cGAMP production in 293T cells 24h later after cGAS^R241E^ / cGAS^Dead^ LNPs treatment was measured by ELISA. (c) mutuDC1940 cell line was treated with cGAS^R241E^ / cGAS^Dead^ mRNA LNPs for 24h. CD40 and CD86 expression was detected with flow cytometry. (d) IFN-β production in wild type (WT) and STING knockout (STINGKO) mutuDC1940 after cGAS^R241E^ / cGAS^Dead^ mRNA treatment for 24h was measured with cytokine bead array (CBA). (e, f) B16 tumor-bearing mice were intratumorally treated with cGAS^R241E^ LNP. Tumor tissue and blood were collected at 6, 12, and 24h. cGAMP level was determined with ELISA (e). IFN-β and TNF cytokines were detected with CBA (f). (g, h) B16 tumor-bearing mice were intratumorally treated with cGAS^R241E^ LNP. cDC1 activation (g) and lymphocyte activation (h) were evaluated 16h later. (i) WT / STING knockout tumor-bearing mice were intratumorally treated with cGAS^R241E^ or control PBS on days 10 and 13. Tumor growth was measured twice a week. **p<0.01, ***p<0.001, ****p<0.0001.

In 293T cells, cGAS^R241E^ mRNA-LNPs induced robust intracellular cGAMP production, whereas cGAS^Dead^ LNPs did not generate detectable cGAMPs (**Fig. 3b**), confirming that cGAMP synthesis depends on intact cGAS catalytic activity. To assess DC activation, we treated the conventional type 1 DC (cDC1) cell line mutuDC1940^34^ and observed significant upregulation of CD40 and CD86 following cGAS ^R241E^ LNP treatment (**Fig. 3c**). Consistently, cGAS ^R241E^ LNP induced robust IFN-β production that plays a pivotal role in enhancing antigen cross-presentation by cDC1^35, 36^, whereas the cGAS^Dead^ LNP had no effect. Notably, IFN-β induction was completely abolished in STING-deficient mutuDC1940 cells, confirming that inflammatory signaling is STING dependent rather than attributable to the lipid carrier or mRNA backbone (**Fig. 3d**).

To determine whether DC activation was driven by unleashed cGAS catalytic activity rather than protein overexpression, we compared cGAS^R241E^ with wild-type cGAS (cGAS^WT^) LNPs. Although cGAS^WT^ moderately increased CD80, CD86 and I-A/I-E expression, cGAS^R241E^ induced markedly stronger upregulation of these activation markers (**Extended Data Fig. 4d-f**). Similarly, only cGAS^R241E^ triggered robust production of downstream cytokines, including IFN-β, TNF and CXCL10, indicating that unleashed catalytic activity is responsible for DC activation (**Extended Data Fig. 4g-i**).

We next examined the *in vivo* uptake of cGAS^R241E^ mRNA-LNPs using DiD-labeled nanoparticles and found the LNPs were efficiently taken up by tumor-infiltrating immune cells, with myeloid cells, particularly macrophages, exhibiting the highest level of uptake (**Extended Data Fig. 4j**). We then evaluated the kinetics of cGAMP production following intratumoral administration of cGAS^R241E^ mRNA LNPs in B16F10 melanoma-bearing mice. cGAMP levels increased rapidly within 6 hours and remained elevated at 24 hours in the tumor, whereas minimal cGAMP was detected in the circulation, consistent with sustained intracellular production and local retention (**Fig. 3e**). Induction of inflammatory cytokines such as IFN-β and TNF followed similar kinetics (**Fig. 3f**), indicating localized innate immune activation within the TME.

We further examined the activation of immune cell subsets within the tumor. cGAS^R241E^ LNP treatment induced marked upregulation of CD40, CD80 and CD86 on intratumoral cDC1cells, a DC subset specialized for cross-presentation of tumor antigens (**Fig. 3g**). In parallel, increased CD69 expression was observed on tumor-infiltrating T cells, NK cells, and NKT cells, indicating activation of both innate and adaptive immune compartments (**Fig. 3h**). Finally, cGAS^R241E^ LNPs substantially inhibited tumor growth in wild-type hosts but not STING-deficient hosts (**Fig. 3i**), demonstrating that therapeutic effect depends on host STING signaling. Collectively, these results establish cGAS^R241E^ LNPs as a potent and localized inducer of STING-dependent innate immune activation *in vivo*.

### Membrane-tethered IL-12 mRNA LNPs restrict cytokine activity to the TME

Our initial functional screening identified IL-12 as a candidate to synergize with STING activation (**Fig. 1d**). Given the severe toxicity associated with soluble IL-12 (sIL-12) administration^26, 27^, we designed a membraned-tethered IL-12 construct to restrict its activity locally. To compare the efficacy and safety of sIL-12 with membrane-tethered IL-12, we generated mRNA constructs encoding either sIL-12 or IL-12 fused to the CD8 transmembrane domain (IL-12TM) (**Fig. 4a**). Surface staining of 293T cells treated with IL-12 mRNA LNPs confirmed high membrane expression of IL-12 in the IL-12TM group but not in the sIL-12 group (**Fig. 4b**). Conversely, high levels of soluble IL-12 were detected in the supernatant of sIL-12-treated cells but not in IL-12TM treated cells (**Fig. 4c**). Functional assessment using HEK-Blue IL-12 reporter cells revealed strong STAT4 activation in response to supernatants from sIL-12-treated cells, whereas minimal reporter activity was observed with supernatants from IL-12TM-treated cells (**Fig. 4d**), indicating negligible cytokine shedding from membrane-tethered format.

**Fig. 4.**
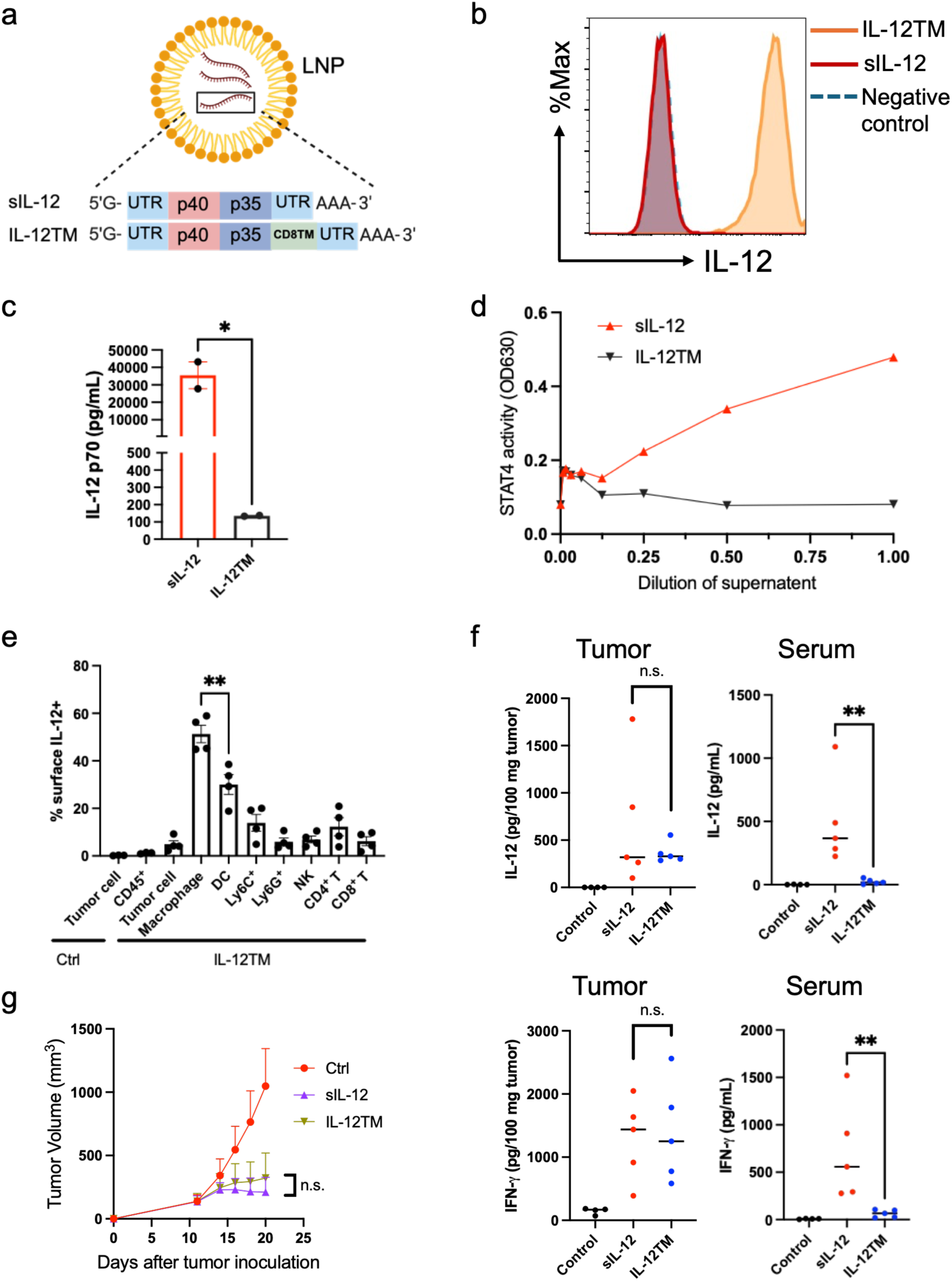
Membrane-tethered IL-12 mRNA LNPs restrict cytokine activity to the TME. (a) Design of soluble IL-12 (sIL-12) and IL-12 with CD8 transmembrane domain (IL-12TM) mRNA constructs. (b-d) 293T cells were treated with sIL-12 and IL-12TM mRNA LNPs for 24h. Surface expression of IL-12 was detected with flow cytometry (b). Supernatant IL-12 level was measured with IL-12 CBA (c). Bioactivity of supernatants was measured with IL-12 reporter cell line (d). (e) IL-12 surface expression on tumor-infiltrating immune cells after intratumoral PBS (Ctrl) or IL-12TM treatment. (f) IL-12 and IFN-γ level in B16 tumor and serum was measured after intratumoral treatment of sIL-12 or IL-12TM mRNA LNPs. (g) B16 tumor bearing mice were intratumorally treated with equal amount of sIL-12 or IL-12TM mRNA LNPs on day 11. Tumor growth was monitored. n.s. p>0.05, *p<0.05, **p<0.01.

We next examined the *in vivo* distribution of IL-12TM following intratumoral delivery. Similar with the pattern of cGAS^R241E^ LNP uptake *in vivo* (**Extended Data Fig. 4j**), immune cells exhibited high surface expression of IL-12, with myeloid populations, particularly macrophages, showing the highest levels of IL-12 expression (**Fig. 4e**). We then compared intratumoral and systemic cytokine levels following administration of equivalent doses of sIL-12 or IL-12TM LNPs. While comparable levels of IL-12 and IFN-γ were detected in the TME, serum concentrations of both cytokines were significantly elevated in the sIL-12 group but remained low in the IL-12TM group, indicating reduced systemic exposure with the membrane-tethered format (**Fig. 4f**). Notably, despite these differences in systemic cytokine levels, both treatments achieved comparable antitumor efficacy (**Fig. 4g**), demonstrating that IL-12TM could deliver therapeutic benefit while mitigating systemic inflammation.

### Combined cGAS^R241E^ and IL-12TM mRNA LNP treatment controls local and distal tumors and reprograms tumor-infiltrating T cells

We employed a B16F10 two-tumor model to evaluate the antitumor efficacy of combined cGAS^R241E^ and IL-12TM mRNA LNPs. Intratumoral administration of either cGAS^R241E^ or IL-12TM LNPs alone inhibited growth of treated tumor but exerted only modest effects on untreated distal tumors. In contrast, Combination treatment (Combo) markedly suppressed tumor growth at both sites (**Fig. 5a**). To assess whether these effects extended to other tumor models, we evaluated Combo in CT26 and TC-1 models and similarly observed enhanced control of both treated and untreated distal tumors (**Fig. 5b and Extended Data Fig. 5a**). Notably, TC-1 is resistant to immune check point blockades (**Extended Data Fig. 5b**).

**Fig. 5.**
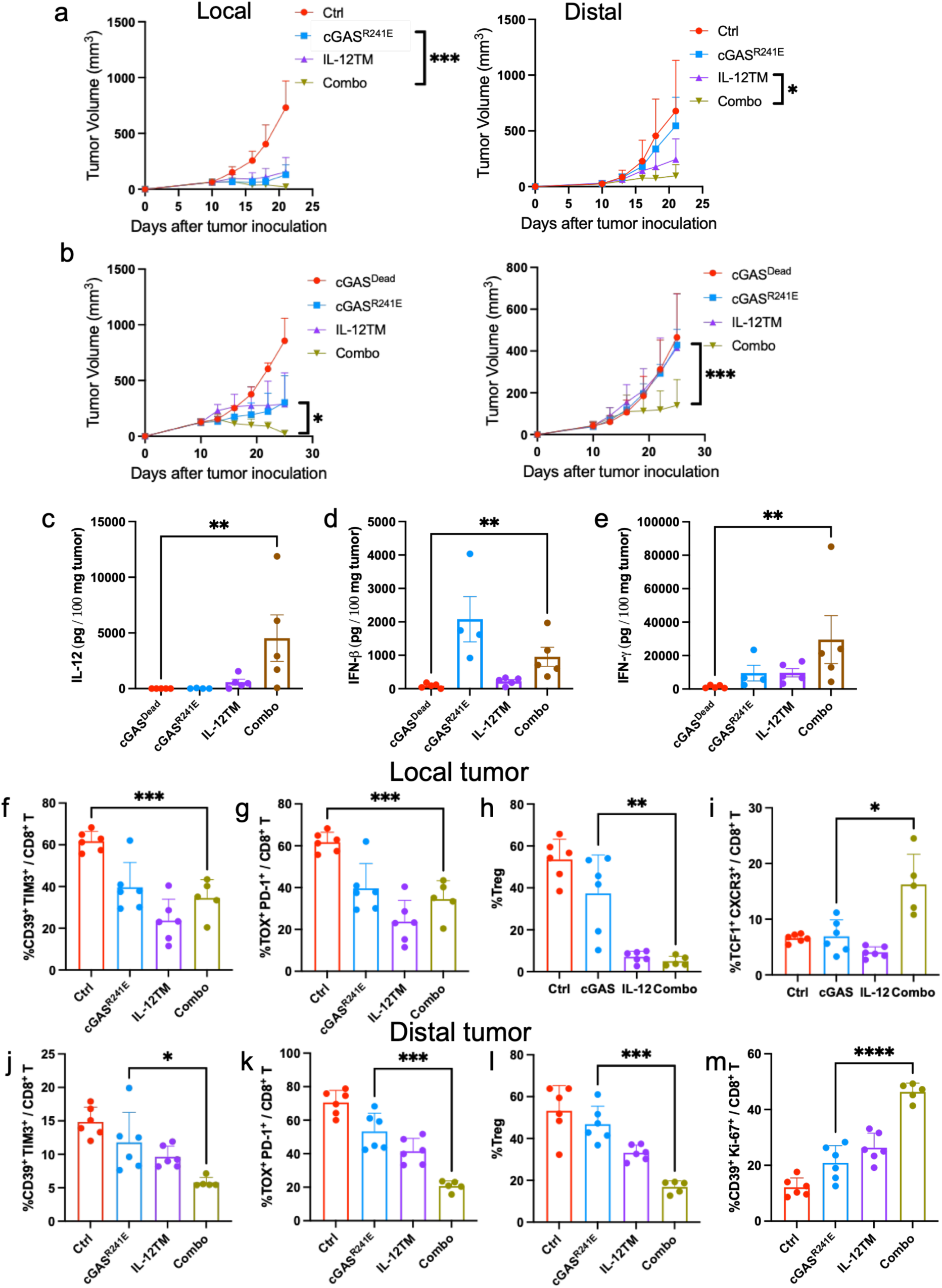
Combined cGAS^R242E^ and IL-12TM mRNA LNP treatment controls local and distal tumors and reprograms tumor-infiltrating T cells. (a) B16 tumor-bearing mice were treated with cGAS^Dead^ control, cGAS^R241E^, IL-12TM, or combination of the two (Combo) on days 10 and 13. Tumor growth was monitored bilaterally. (b) CT26 tumor-bearing mice were treated with cGAS^Dead^ control, cGAS^R241E^, IL-12TM, or combination of the two (Combo) on days 10, 13, 16 and 19. Tumor growth was monitored bilaterally. (c-e) B16 tumor-bearing mice were treated with cGAS^Dead^ control, cGAS^R241E^, IL-12TM, or Combo. Tumor tissues were analyzed for IL-12 (c), IFN-β (d) and IFN-γ (e) production by CBA 16h later. (f-m) B16 tumor-bearing mice were treated with cGAS^R241E^, IL-12TM or Combo. Frequency of CD39^+^ TIM3^+^ CD8^+^ T cells (f), TOX^+^ PD1^+^ CD8^+^ T cells (g), Tregs (h) and TCF1^+^CXCR3^+^ CD8^+^ T cells (i) from local tumors. Frequency of CD39^+^ TIM3^+^ CD8^+^ T cells (j), TOX^+^ PD1^+^ CD8^+^ T cells (k), Tregs (l) and CD39^+^ Ki67^+^ CD8^+^ T cells (m) from distal tumors. *p<0.05, **p<0.01, ***p<0.001, ****p<0.0001.

We then evaluated the treatment-associated toxicity. No significant body weight loss was observed after Combo treatment (**Extended Data Fig. 5c**). We further measured serum creatinine and alanine aminotransferase levels which indicate kidney and liver function respectively, and found no significant changes among treatment groups (**Extended Data Fig. 5d)**. Similarly, histological examination revealed no apparent tissue damage (**Extended Data Fig. 5e**). Together, these findings demonstrate that combining cGAS– STING activation with IL-12 signaling through mRNA-LNP delivery enhances local and distal tumor control without inducing detectable systemic toxicity.

To define how Combo treatment reshapes the TME, we first analyzed intratumoral cytokine profiles. Combo therapy markedly increased intratumoral levels of IL-12 and IFN-γ, while the main contribution of IFN-β is from cGAS^R241E^ treatment (**Fig. 5c-e**). Flow cytometric analysis of tumors revealed pronounced remodeling of immune cell populations. In local tumors, Combo therapy reduced the frequencies of T cells expressing dysfunction-associated markers, including CD39, TIM-3, PD-1, and TOX, decreased Treg abundance, and expanded T cell populations expressing TCF1 and CXCR3 (**Fig. 5f-i**). Similar reductions in dysfunction-associated T cell populations and Tregs were observed in untreated distal tumors, where the effects of combination treatment were more pronounced (**Fig. 5j-l**). Combo therapy also enhanced T cell proliferation while reducing the proportion of cells exhibiting a dysfunction-associated phenotype (**Fig. 5m**). These findings are consistent with the transcriptional remodeling identified by scRNA-seq (**Fig. 2**). Collectively, these findings demonstrate that combined cGAS^R241E^ and IL-12TM mRNA-LNP therapy induces coordinated cytokine responses and reprograms the TME at both local and systemic sites to promote systemic tumor control.

### Control of distal tumors by combined cGAS^R241E^ and IL-12TM LNP therapy requires CD8^+^ T cells and IFN-γ signaling

To define the immune mechanisms underlying systemic tumor control, we evaluated the Combo therapy in *Rag1^−/−^*mice. While suppression of locally treated tumors was partially preserved, the anti-tumor effect against distal tumors was completely abrogated, indicating an essential role for adaptive immunity in mediating systemic tumor control (**Fig. 6a**). Consistently, depletion of CD8⁺ T cells completely abolished distal tumor regression, while local tumor suppression remained largely intact (**Fig. 6b**), suggesting that local tumor control may involve direct STING-mediated tumor killing. Additional depletion of CD4⁺ T cells, NK cells, or neutrophils did not significantly alter tumor control (**Extended Data Fig. 6a, b**). Because BATF3-dependent DCs are classically required for tumor antigen cross-presentation, we next evaluated Combo therapy in *Batf3^⁻/⁻^* mice. Remarkably, the Combo treatment retained its ability to suppress distal tumor growth in the absence of BATF3-dependent DCs (**Fig. 6c**), suggesting that this DC subset is not strictly required for the systemic antitumor activity of the combination. Combo efficacy was partially dependent on host STING signaling (**Extended Data Fig. 6c**), consistent with the contribution from the cGAS^R241E^ mRNA-LNP component.

**Fig. 6.**
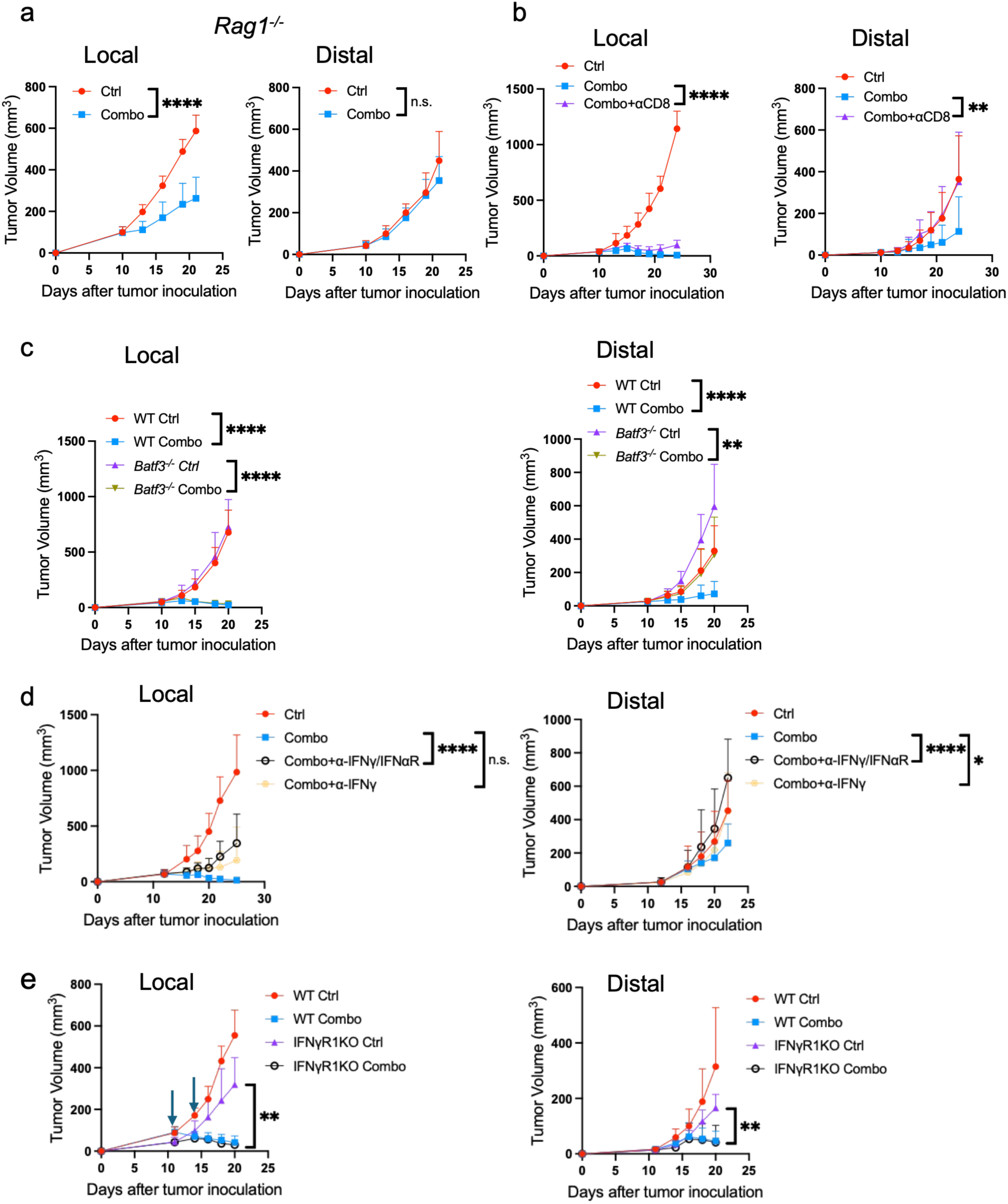
Control of distal tumors by combined cGAS^R241E^ and IL-12TM LNP therapy requires CD8^+^ T cells and IFN-γ signaling. (a) *Rag1^−/−^*mice bearing B16 tumors were treated with Combo LNPs. Tumor growth was monitored bilaterally. (b) B16 tumor bearing mice were treated with Combo LNPs. CD8 depletion antibodies were intraperitoneally injected twice a week starting on one day before treatment. Tumor growth was monitored bilaterally. (c) *Batf3^−/−^*mice bearing B16 tumors were treated with Combo LNPs. (d) B16 tumor bearing mice were treated with Combo LNPs on days 12 and 15. Anti-IFN-γ and anti-IFNαR antibodies were intraperitoneally injected twice a week starting on one day before treatment. Tumor growth was monitored bilaterally. (e) Wild type B16 and IFNγR1KO B16 tumor bearing mice were treated with Combo LNPs. Tumor growth was monitored bilaterally. n.s. p>0.05, *p<0.05, **p<0.01, ****p<0.0001.

Given the robust induction of both type I and type II IFNs following Combo treatment, we next interrogated their functional relevance. Blockade of IFN-I signaling alone did not significantly affect the anti-tumor efficacy. In contrast, inhibition of IFN-γ signaling completely abolished the suppression of distal tumor growth, while combined blockade of IFN-I and IFN-γ pathways further compromised the tumor control at both local and distal sites (**Fig. 6d** and **Extended Data Fig. 6d**). Consistent with these *in vivo* findings, ex vivo stimulation of splenocytes from IFN-γ YFP reporter mice revealed increased IFN-γ production by CD8^+^ T cells, NK cells and NKT cells following Combo treatment (**Extended Data Fig. 6e**). We further evaluated whether the effects of IFN-γ is through direct tumor killing by knocking out IFNγR on tumor cells (**Extended Data Fig. 6f**). Combo treatment retained its antitumor activity against both treated and distal IFNγR-deficient tumors, suggesting that tumor cell–intrinsic IFN-γ signaling is dispensable for therapeutic efficacy. IFN-γ may instead act through host cells, potentially including Treg, which has been suggested by previous studies^37, 38^.

### Combo treatment synergizes with PD-L1 blockade to enhance tumor control and survival

We observed that both cGAS^R241E^ LNP and IL-12TM treatment induced marked upregulation of PD-L1 expression on macrophages and DCs in both local and distal tumors, suggesting the emergence of adaptive immune resistance (**Fig. 7a, b**). To overcome this resistance, we combined Combo treatment with PD-L1 blockade. This triple combination significantly enhanced distal tumor control and prolonged overall survival compared with either treatment alone (**Fig. 7c, d, Extended Data Fig. 7a**). Notably, PD-L1 blockade alone showed minimal efficacy in this two-tumor model, indicating that Combo therapy sensitizes otherwise PD-L1 blockade-refractory tumors to immune checkpoint inhibition. A similar survival benefit was observed in CT26 tumor model (**Extended Data Fig. 7b, c**), indicating that the benefit of combing Combo with PD-L1 blockade was not restricted to a single tumor model.

**Fig. 7.**
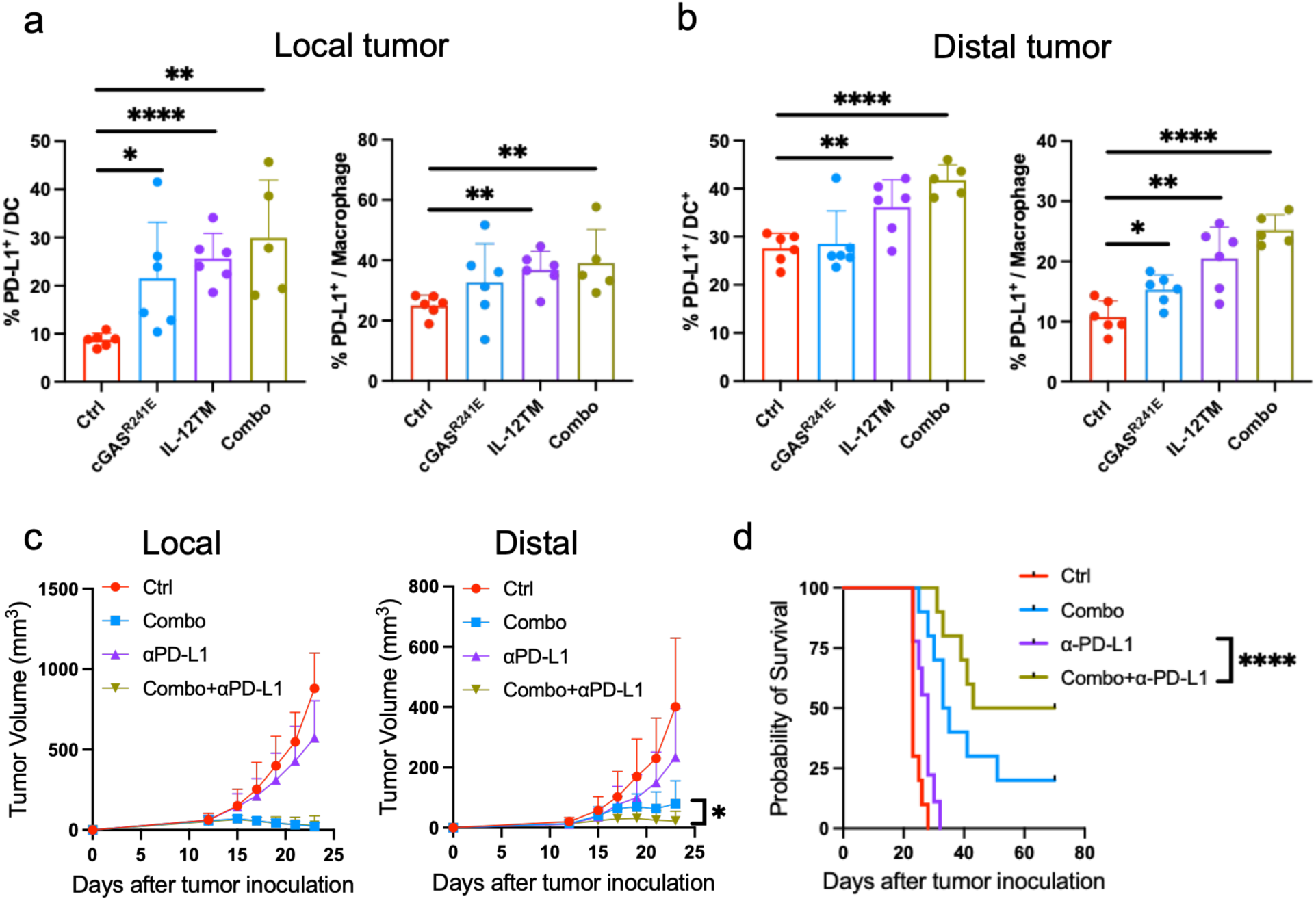
Combo treatment synergizes with PD-L1 blockade to enhance tumor control and survival. (a, b) B16 tumor-bearing mice were treated with PBS, cGAS^R241E^, IL-12TM or Combo. DC and macrophages from local (a) and distal tumors (b) were analyzed for expression of PD-L1. (c, d) B16 tumor-bearing mice were treated with PBS, Combo, αPD-L1 or the triple combination (Combo+αPD-L1) on days 12 and 15. Tumor growth was monitored bilaterally (c). (d) Overall survival capacity was measured (data pooled from two experiments). *p<0.05, **p<0.01, ***p<0.001, ****p<0.0001.

## Discussion

Local cancer therapies frequently fail to elicit durable systemic antitumor immunity capable of controlling metastatic disease. Although current approaches such as STING agonists and cytokine-based therapies have shown promise, their clinical utility has been limited by transient efficacy and dose-limiting systemic toxicity. Here we demonstrate that combined delivery of the autoactivated cGAS^R241E^ mutant and membrane-tethered IL-12 via mRNA LNPs induces potent anti-tumor immunity against both local and distal tumors across multiple preclinical models. The cGAS^R241E^ mRNA LNPs drive robust, localized cytokine production and immune activation within the TME without triggering systemic inflammation. In parallel, IL-12TM mRNA LNPs limit circulating IL-12 and IFN-γ while preserving local cytokine activity. Together, these complementary mechanisms inhibit Tregs and reprogram intratumoral T cells toward less dysfunctional effector states, enabling synergistic control of distal tumors in a CD8⁺ T cell- and IFN-γ–dependent manner and enhancing responsiveness to PD-L1 blockade.

Despite compelling preclinical efficacy, clinical translation of STING pathway activation has been hindered by pharmacokinetic and safety limitations of cyclic dinucleotide agonists, including poor permeability, short half-life and systemic toxicity^13, 39^. The cGAS^R241E^ mRNA LNP strategy addresses several of these limitations. Intracellular expression of cGAS^R241E^ enables sustained cGAMP production within tumors, overcoming the delivery and permeability limitations associated with exogenous agonists. In addition, cell-intrinsic cGAMP generation promotes localized immune activation while minimizing systemic exposure. The mRNA-LNP platform may further contribute to immune modulation within the TME. Collectively, these features position cGAS^R241E^ mRNA LNPs as an alternative approach to conventional STING agonists with improved pharmacologic properties.

While cGAS^R241E^ mRNA LNPs induce a broad inflammatory response, they have limited effect on IL-12, a key cytokine that amplifies IFN-γ signaling and sustains cytotoxic T cell function^40^. Previous work demonstrated that combining the STING agonist MSA-2 with mRNA LNPs encoding secreted IL-12 enhanced T cell function and antitumor efficacy^41^. Our approach differs by coupling intracellular cGAMP production driven by mRNA-encoded cGAS^R241E^ with membrane-tethered IL-12, which is designed to localize cytokine activity within the tumor microenvironment and limit systemic exposure. Combing IL-12TM with cGAS^R241E^ mRNA LNPs enhanced IFN-γ production, reduced Tregs, and shifted tumor-infiltrating T cells toward less dysfunctional, more activated effector states, enabling effective control of distal tumors. Notably, the antitumor effects of the combination depend on IFN-γ but not on tumor cell–intrinsic IFN-γ signaling, suggesting that IFN-γ acts through host cells. Modulation of Treg function may contribute to this response, consistent with previous reports that IFN-γ can induce Treg fragility and that IL-12 can suppress Treg-mediated immune regulation through IFN-γ-dependent mechanisms^37, 38^.

Mechanistically, tumor control induced by the combination therapy reflects coordinated contributions from innate and adaptive immunity. Partial preservation of local tumor suppression in *Rag1^⁻/⁻^* mice suggests that innate immune activation contributes to local tumor regression, consistent with robust STING-dependent signaling. In contrast, distal tumor control was strictly dependent on CD8⁺ T cell and IFN-γ signaling, underscoring their indispensable roles in systemic antitumor immunity. Interestingly, this CD8⁺ T cell-dependent response was maintained in *Batf3^−/−^*mice, indicating that combined activation of STING and IL-12 pathways can bypass canonical BATF3-dependent cDC1-mediated cross-priming. A similar mechanism has recently been observed in the context of mRNA-LNP vaccination^42^. This is particularly intriguing given the scarcity of cDC1s in many tumors, which represents a major barrier to effective antigen presentation and immunotherapy responsiveness^43^.

In summary, we establish an mRNA-LNP–based immunotherapy strategy that couples cGAS-STING–driven innate immune activation with localized IL-12 signaling to elicit systemic antitumor immunity. This approach induces CD8⁺ T cell-and IFN-γ–dependent tumor control while limiting systemic cytokine exposure and enhances responsiveness to PD-L1 blockade. Built on clinically validated mRNA-LNP delivery technology with clear translational feasibility, this strategy provides a clinically actionable framework for coordinating innate and adaptive immunity to achieve durable systemic tumor control.

## Acknowledgments

Z.J.C was funded by NIH (U54CA244719, R01CA299257), Welch Foundation (I-1389), and Cancer Grand Challenge from CRUK (CGCATF-2021/100023) and NCI (OT2CA278683). A.Z was funded by Lymphoma Research Foundation, Breast Cancer Alliance and Susan G. Komen Foundation. Z.J.C is an investigator of the HHMI. Y.F. was supported by the NCI Predoctoral to Postdoctoral Fellow Transition Award (F99CA274647/K00CA274647).

## Author Contributions

Conceptualization, A.Z., Y.F. L.S. and Z.J.C.; Methodology, A.Z., Y.F., R.P., M.Y., Z.H.C., T.L., J.T.; Investigation, A.Z. L.S. and Y.F.; Writing - Original Draft, A.Z.; Writing-Review & Editing, Y.F. and Z.J.C.; Funding Acquisition, Z.J.C., A.Z. and Y.F.; Supervision, A.Z. and Z.J.C.

## Methods

Key reagents and assays details were provided in **Supplementary Table**.

### Experimental Model

#### Mice

C57BL/6J, *Tmem173*^−/−^, *Batf3^−/−^*, *Rag1^−/−^* and IFN-γ reporter mice (*Ifng^tm3.1Lky^*/J) in the C57BL/6J background were purchased from the Jackson Laboratory. All mice were maintained under specific pathogen-free conditions. Animal care and experiments were carried out under institutional and National Institutes of Health protocol and guidelines. This study has been approved by the Institutional Animal Care and Use Committee of the University of Texas Southwestern Medical Center.

#### Cell lines and reagents

B16, TC-1, CT26 and 293T cells were purchased from ATCC. HEK-Blue 293 IL-12 reporter cells are purchased from InvivoGen. The above cells were cultured in Dulbecco’s modified Eagle’s medium supplemented with 10% heat-inactivated fetal bovine serum (FBS), 100 U/mL penicillin, and 100 U/mL streptomycin. MutuDC1940 were from Applied Biological Materials HEK Blue 293 reporter cell and were cultured in IMDM supplemented with 10% FBS, 50 µM β-mercaptoethanol,10 mM HEPES and 1% Penicillin/Streptomycin Solution. All cell lines were cultured under 5% CO_2_ at 37 °C and were routinely tested for mycoplasma contamination with PCR.

Anti-CD4 (GK1.5), anti-CD8 (Lyt3.2), anti-NK1.1 (PK136), anti-Ly6C (Monts1), anti-Ly6G (Gr-1), anti-IFN-γ (XMG1.2), anti-CTLA-4 (9D9) and anti-IFNαR-1(MAR1-5A3) antibodies were purchased from BioXCell. Anti-PD-L1 (atezolizumab) were kindly provided by UT Southwestern Simmons Cancer Center Pharmacy.

### Method Details

#### mRNA LNP design, synthesis and characterization

Mouse cGAS^R241E^, cGAS^Dead^ and cGAS^WT^ sequences were cloned via Gibson Assembly into an mRNA expression vector containing optimized 5′ and 3′untranslated regions (UTRs) and a poly (A) tail. For soluble mIL-12 and IL-12TM, an (G_4_S)_3_ linker was inserted between IL12b and IL12a sequences. For IL-12TM, the C terminus of IL12b-IL12a construct was linked via additional (G_4_S)_3_ linker to mouse CD8 transmembrane domain. mRNAs were synthesized in vitro using T7 RNA polymerase.

For LNP formulations, lipids were dissolved in ethanol at molar ratios of 50:10:38.5:1.5 (SM102: DSPC: cholesterol: PEG-2000). The lipid mixture was mixed with mRNA diluted in 50 mM sodium acetate buffer (pH 4) using a microfluidic mixer (Precision Nanosystems) at a ratio of 3:1 (aqueous: ethanol). mRNA LNPs were dialyzed in PBS in dialysis cassettes for overnight and concentrated using Amicon Ultra centrifugal filters (EMD Millipore). Particle size and zeta potential of the LNPs were measured by dynamic light scattering instrument (Anton Paar). The mRNA encapsulation efficiency was quantified using the Quant-iT RiboGreen assay.

#### cGAMP ELISA and cytokine bead array

B16 tumors were harvested, and 20-30 mg tumor tissues were homogenized with a tissue homogenizer (MP FastPrep-24 5G) in tissue protein extraction reagent with protease inhibitors. Supernatants were collected, and analyzed for cGAMP level using ELISA (Cayman) and for inflammatory cytokines with LEGENDplex™ Mouse Anti-Virus Response Panel (Biolegend).

#### Tumor growth and treatment

A total of 5×10^5^ B16, 2.5×10^5^ TC-1 or 5×10^5^ CT-26 cells were inoculated subcutaneously into right dorsal flanks of the mice in 100 μL PBS on day 0. In two-tumor model, the same number of tumor cells were inoculated subcutaneously into left dorsal flanks on day 3. For initial cytokine screening experiment, 5×10^5^ or 2×10^5^ B16 tumor cells were inoculated on right or left dorsal flanks of mice on day 0. Tumor-bearing mice were randomly grouped into treatment groups when right side of tumors reached ∼100 mm^3^. All treatments were administrated intratumorally, cGAMP (10 µg), sumIL-2 (500 ng), IL-15 (rIL-15: 80 ng, rIL-15Rα: 480 ng), IL-21 (50 ng) and IL-12-Fc (50 ng, mEq 30ng) cytokines were injected on days 10,14 and 18; cGAS^Dead^ (5 µg), cGAS^R241E^ ( 2.5 or 5 µg) and sIL-12/IL-12TM mRNA LNP ( 50-250 ng) were injected on days 10 and 13. IL-15 and sIL-15Ra-Fc were incubated together for 20 min at 37 °C before injection^44^. Cytokines were prepared in PBS with 1% mouse serum. For combination of cytokines with cGAMP treatment, cytokine and cGAMP were mixed immediately before injection to avoid instability of cGAMP in serum.

For immune cell depletion or blockade studies, 200 µg of antibodies were intraperitoneally injected one day before treatment initiation and then twice a week for 2 weeks. For PD-L1 blockade combination therapy in B16 model, 200 μg PD-L1 was intraperitoneally injected on days 11 and 14. Tumor volumes were measured by the length (a), width (b) and height (h) and calculated as tumor volume = abh/2.

#### Single cell RNA sequencing

Two B16 tumor models were established as described above. Local (right) tumors were treated with cGAMP (10 µg) and IL-12 (30 ng) on days 9, 13, and 17. For scRNA-seq, samples from five control mice or eight cGAMP+IL-12-treated mice were pooled within each treatment condition and tissue compartment before library preparation. Mice were sacrificed on day 21, and both local and distal tumors were harvested and processed into single-cell suspensions. CD45-positive immune cells were enriched using the Mouse TIL (CD45) Positive Selection Kit, followed by staining with CD3, NK1.1 and CD11c and sorting on an Aria I flow sorter (BD Biosciences) to enrich T cells, NK cells, and dendritic cells. Collected cells were processed for single-cell RNA sequencing. Libraries for gene expression (GEX) and TCR V(D)J sequencing were prepared for each sample following the 10x Genomics User Guide (CG000208 Rev G) and sequenced by Novogene.

#### Quality control for GEX analysis

Single-cell RNA-seq data were analyzed using Seurat (v4.1.0) in R. Low-quality cells were excluded based on high mitochondrial gene content (percent.mt > 10%) or abnormal feature counts (nFeature_RNA < 500 or > 6,000). Potential doublets were identified using DoubletFinder; clusters in which more than 50% of cells were labeled as doublets, as well as individual doublets, were removed. Remaining singlet cells were integrated using Seurat’s IntegrateData function.

Integrated cells were clustered using FindClusters across multiple resolutions (0.4, 0.6, 0.8, 1.0, and 1.4). A resolution of 1.0 provided clear separation of major immune populations, including CD8 T cells, CD4 T cells, NK cells, myeloid cells, and B cells, based on canonical markers such as Cd3d, Cd4, Cd8b1, Klrb1c, Itgax/Itgam, and Cd19. CD8 T cells and CD4 T cells were subsequently subset for lineage-specific analyses.

#### Single-cell RNA-sequencing analysis

For higher-resolution analysis, CD8⁺ and CD4⁺ T cells were reclustered independently using the first 20 principal components. Clustering was evaluated at resolutions 0.4, 0.6, 0.8, 1.0, and 1.4, and resolution 1.0 was selected based on cluster structure and marker gene expression. Robustness was assessed by tracking overlap of the final annotations across resolutions using precision, recall, and their harmonic mean (cell-overlap F1) and by visualizing cluster relationships with clustree. Alternative t-SNE embeddings were generated from the same PCA representation. UMAP robustness was evaluated across five random seeds and parameter settings spanning n.neighbors = 15-50 and min.dist = 0.1-0.5. Marker expression was quantified from the RNA assay, and transcriptional program scores were calculated using Seurat AddModuleScore. Closely related graph clusters were merged when their principal distinction reflected cell-cycle activity rather than a distinct functional program; the two graph clusters comprising the final C10 dysfunction-associated state were merged on this basis. Differentially expressed genes for each cluster were identified using Seurat’s FindAllMarkers function with only.pos = TRUE, min.pct = 0.5, and logfc.threshold = 0.5. The primary analysis used Seurat v4.1.0; robustness reanalysis was performed with Seurat v5.4.0.

#### GO enrichment analysis

Gene Ontology (GO) enrichment analysis was performed to identify signaling pathways enriched in each cell cluster. Cluster-specific differentially expressed genes were identified using Seurat (v4) as described above. GO enrichment analysis was conducted using the enrichGO function from the clusterProfiler package (ontology = “BP”), with multiple testing correction by the Benjamini–Hochberg method and significance defined as q value < 0.05. To improve visualization and reduce redundancy among GO terms, enriched GO Biological Process results were simplified based on semantic similarity using the simplify () function (cutoff = 0.7).

#### Flow cytometry analysis

Single-cell suspensions were incubated with anti-FcγIII/II receptor (clone 2.4G2) for 15 mins to block the non-specific binding, followed by staining with fluorophore-conjugated antibodies for 30 mins at 4°C in the dark. Fixable viability Dye live dead blue (eBioscience) was used to exclude the dead cells. Samples were acquired on Cytek Aurora flow cytometer and analyzed by using FlowJo software (BD).

#### Cryo-EM imaging of mRNA LNPs

To prepare cryo-EM grids, 3 μL of cGAS^R241E^ LNP solution was applied to a glow-discharged copper grid with a 2 nm ultra-thin carbon support film (Quantifoil R1.2/1.3, 300 mesh). Blotting was performed for 4 seconds at a blot force of –5, 6 °C, and 100% humidity using an FEI Vitrobot Mark IV. After a 60-second wait, the grid was blotted, rapidly plunged into liquid ethane, and stored in liquid nitrogen until imaging. Cryo-EM data were collected on a Thermo Fisher Glacios electron microscope operated at 200 kV and equipped with a K3 direct electron detector at the UTSW cryo-EM facility. The nominal magnification was 45,000×.

#### Quantification and Statistical Analysis

All the data analyses were performed with GraphPad Prism statistical software and shown as mean ± SEM. P value determined by two-way ANOVA for tumor growth or Log-rank test for survival or unpaired two-tailed t-tests for other analysis. A value of p < 0.05 was considered statistically significant. Because the scRNA-seq libraries were generated from pooled mice, treatment-associated scRNA-seq comparisons were treated descriptively; individual cells were not treated as biological replicates, and no cell-level treatment significance tests were performed.

## Supplemental Information

**Extended Data Fig.1, related to Fig. 1.**
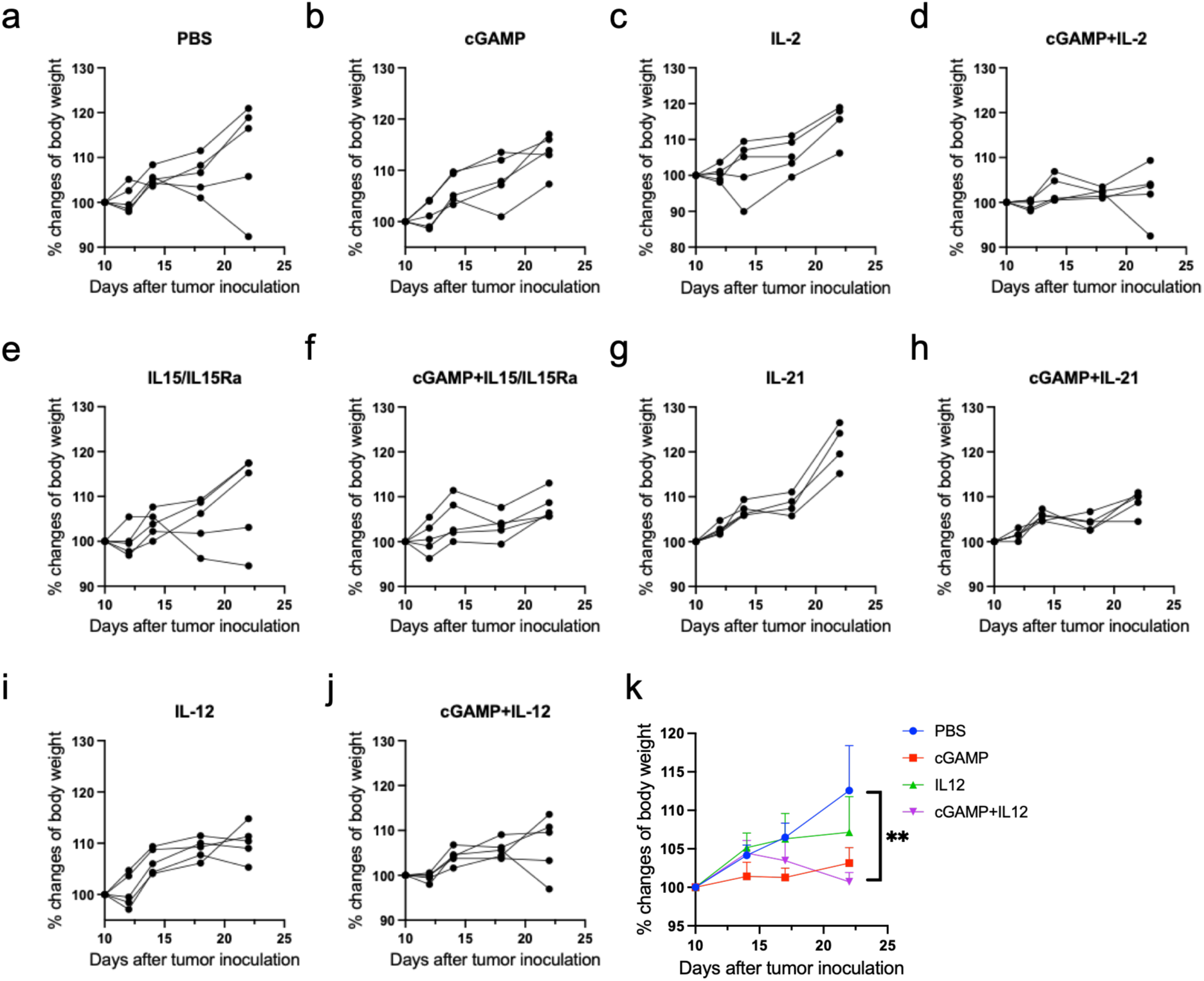
Treatment was shown as in Fig. 1. (a-j) Individual mouse body weight change after each treatment as shown on top of each panel. (k) Pooled mouse body weight changes after PBS, cGAMP, IL-12 or cGAMP plus IL-12 treatment. **p<0.01.

**Extended Data Fig.2, related to Fig. 2.**
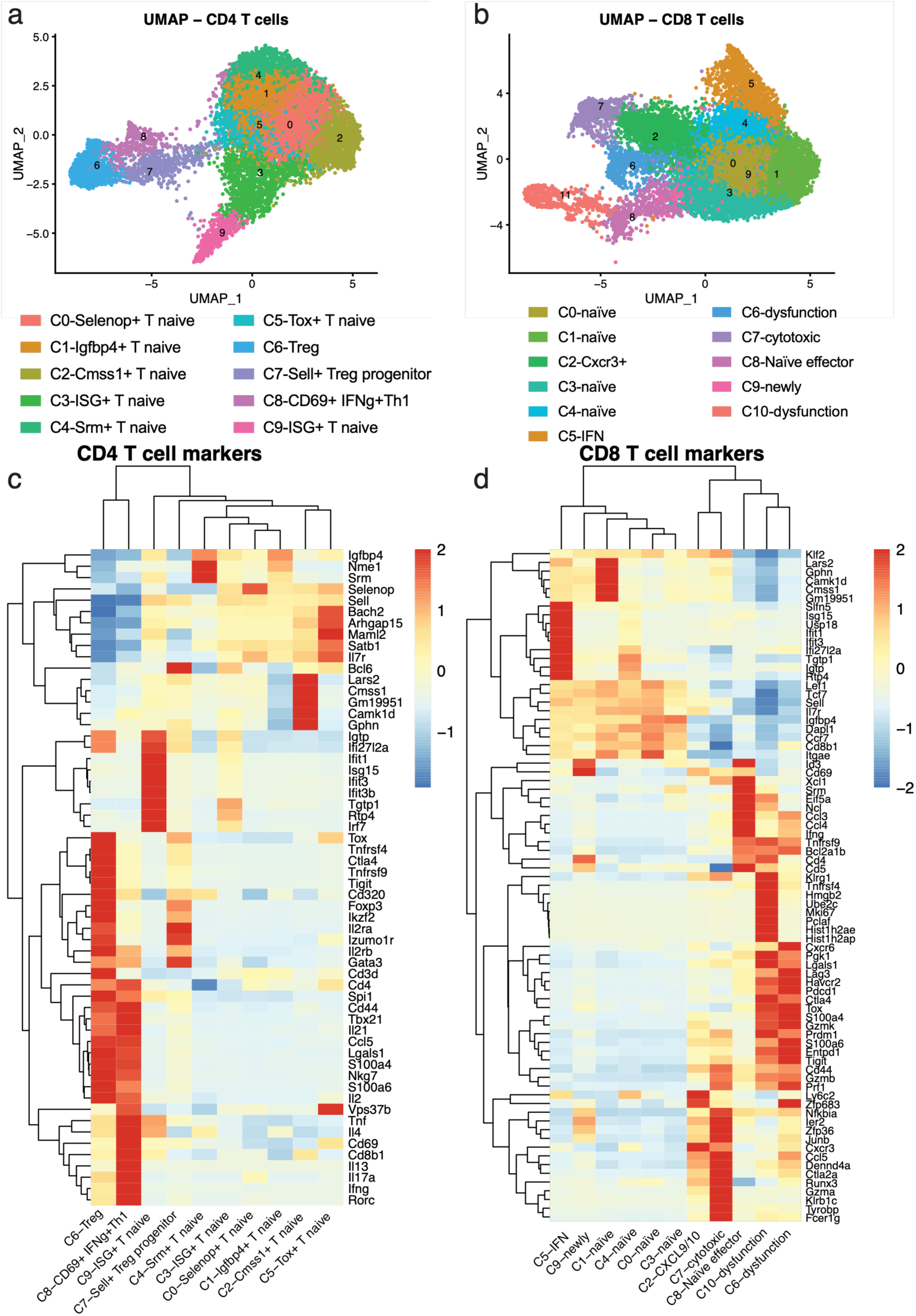
(a, b) UMAP plot showing unsupervised clustering and distribution of total CD4^+^ (a) and CD8^+^ (b) T cells from both local and distal tumors. (c, d) signature genes defining each cluster from CD4^+^ (c) and CD8^+^ (d)T cells were shown.

**Extended Data Fig.3, related to Fig. 3.**
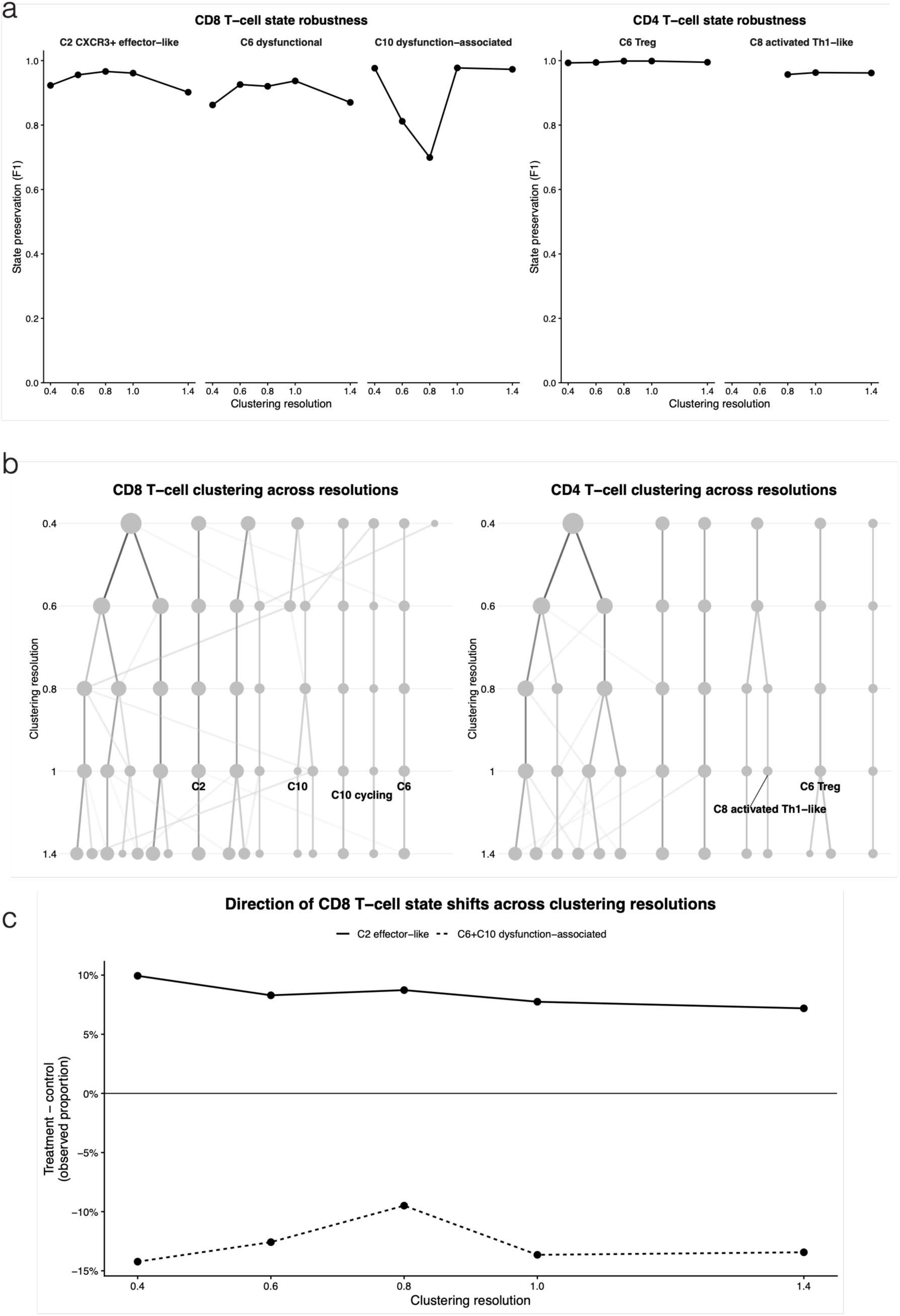
(a) Preservation of key CD8⁺ (left) and CD4⁺ (right) T-cell states across clustering resolutions, quantified by cell-overlap F1 score. (b) Clustree visualization of CD8⁺ and CD4⁺ T-cell clustering across resolutions 0.4-1.4; key states at resolution 1.0 are labeled. (c) Treatment-control difference in observed proportions of the C2 effector-like and combined C6^+^ C10 dysfunction-associated CD8⁺ compartments across resolutions.

**Extended Data Fig.4, related to Fig. 3.**
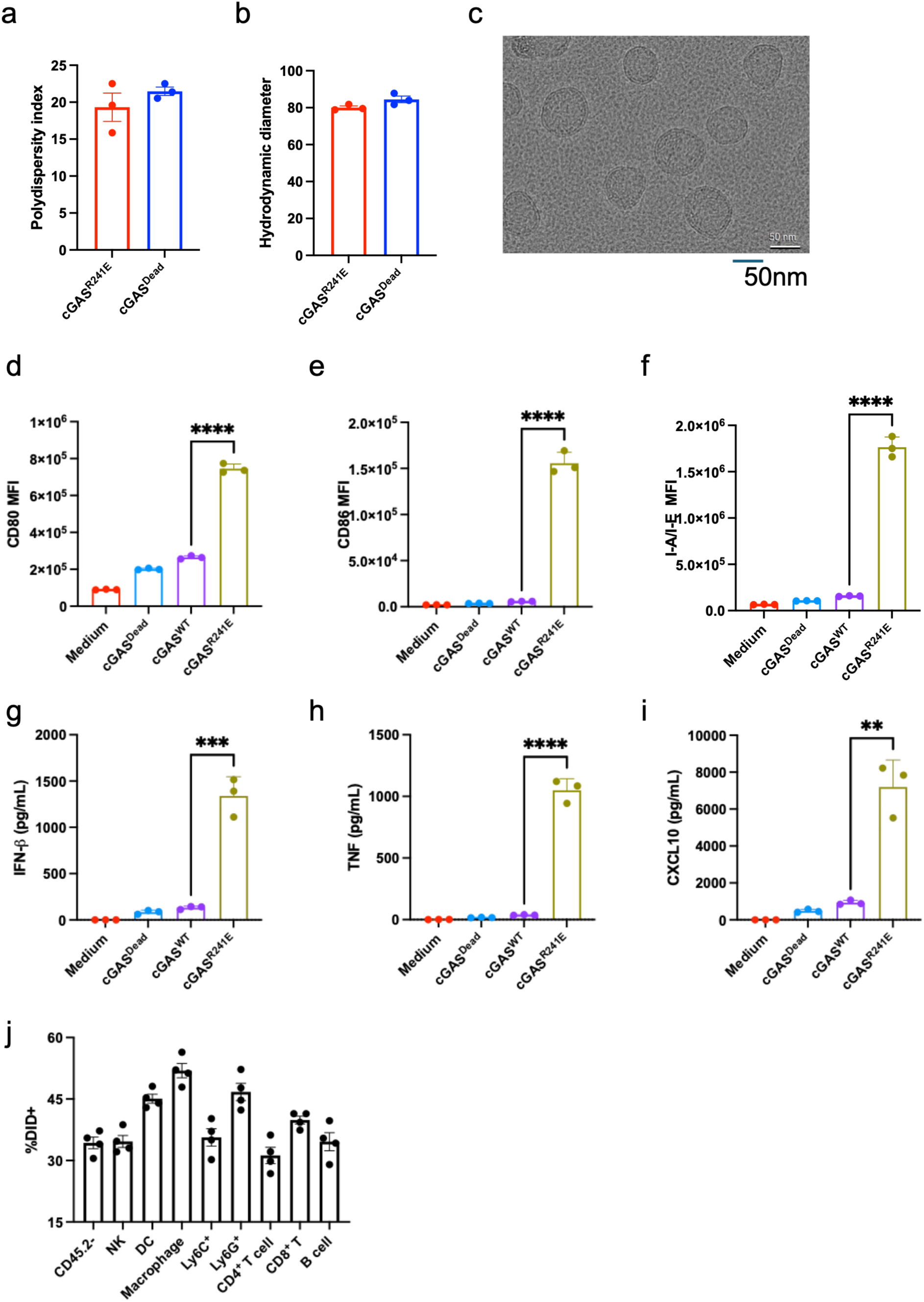
(a, b) Polydispersity index (a) and hydrodynamic diameter (b) of cGAS^R241E^ and cGAS^Dead^ mRNA LNPs measured by dynamic light scattering analysis. (c) Cryo-electron microscopy imaging of cGAS^R241E^ LNPs. Scale bar, 50 nm. (d-i) MutuDC1940 cells were stimulated with cGAS^Dead^, cGAS^WT^ or cGAS^R241E^ LNPs at the presence of 5 μg /mL ApoE for 20hrs. Median fluorescence intensity (MFI) of CD80 (d), CD86 (e) and I-A/I-E (f) measured by flow cytometry were shown. Levels of IFN-β (g), TNF (h) and CXCL10 (i) cytokines in the supernatant were measured by cytokine bead array. (j) cGAS^R241E^ mRNA LNPs were labeled with Did dye and injected into B16-tumor bearing mice. Four hours later, tumors were harvested and Did+ cells were evaluated with flow cytometry. **p<0.01, ***p<0.001, ****p<0.0001

**Extended Data Fig.5, related to Fig. 5.**
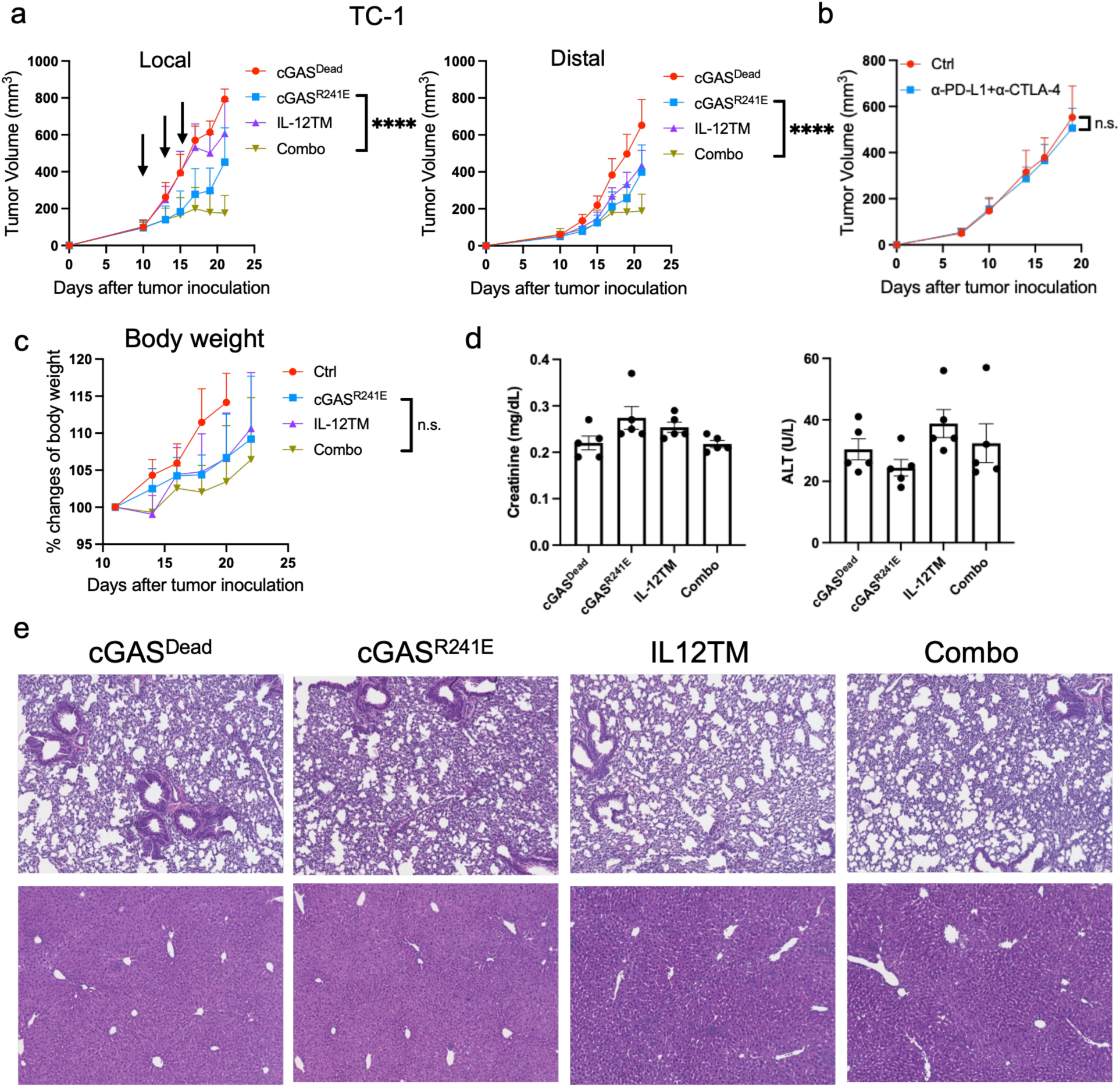
(a) TC-1 tumor-bearing mice were locally treated with cGAS^R241E^, IL-12TM, or Combo. Tumor growth was monitored bilaterally. (b) TC-1 tumor-bearing mice were systemically treated with 200 μg αPD-L1 plus αCTLA-4 on days 7, 10 and 13. Tumor growth was monitored bilaterally. (c) B16 tumor-bearing mice were treated with cGAS^R241E^, IL-12TM or the combination of cGAS^R241E^ and IL-12TM on days 11 and 14. Mice body weight was monitored twice a week. (d, e) B16-tumor bearing mice were treated with cGAS^R241E^ and IL-12TM mRNA LNPs on days 11 and 14. Sera and tissues were collected on day 15. (d) Serum creatinine (left) and alanine aminotransferase (ALT) (right) levels were measured using VITROS chemistry analysis. (e) Representative histological staining of lung (upper panel) and liver tissues (lower panel) after treatments. n.s. p> 0.05, ****p<0.0001.

**Extended Data Fig.6, related to Fig. 6.**
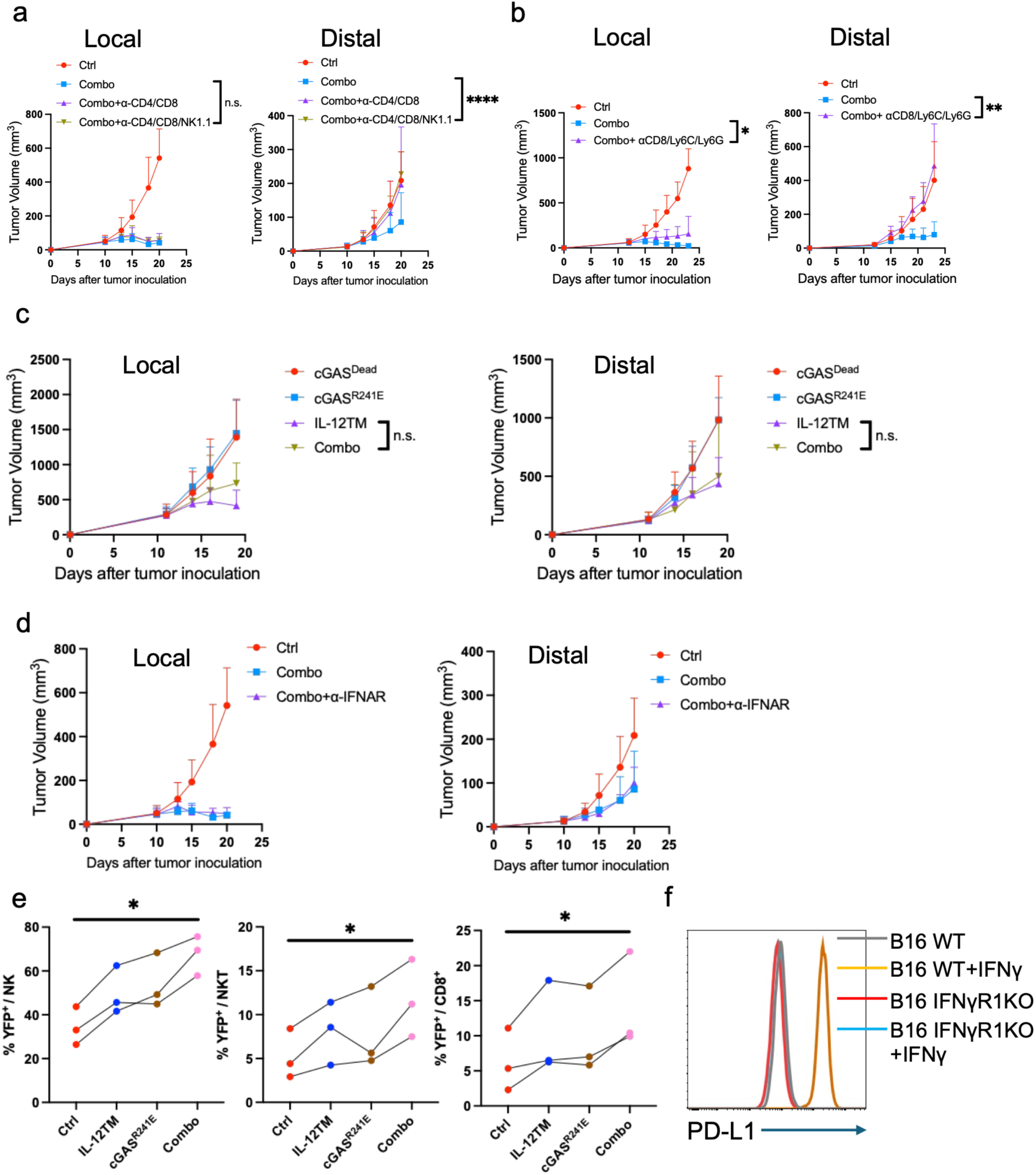
(a, b) B16 tumor bearing mice were treated with Combo LNPs. Anti-CD8/CD4/NK1.1 antibodies (a) or anti-CD8/Ly6C/Ly6G antibodies (b) were intraperitoneally injected twice a week starting on one day before treatment. Tumor growth was monitored bilaterally. (c) STING knockout mice bearing B16 tumors were intratumorally treated with cGAS^R241E^, IL-12TM, or Combo. Tumor growth was monitored bilaterally. (d) B16 tumor bearing mice were treated with Combo LNPs, anti-IFNAR antibodies were intraperitoneally injected twice a week starting on one day before treatment. Tumor growth was monitored bilaterally. (e) Splenocytes from IFN-γ reporter mice were treated with cGAS^R241E^, IL-12TM or Combo for 20h. Percentage of YFP^+^ cells among NK, NKT and CD8^+^ T cells were evaluated. (f) Validation of IFNγR1KO B16 tumor cells. WT or IFNγR1KO B16 tumor cells were treated with 20ng/mL IFN-γ for overnight, PD-L1 expression was measured with flow cytometry. n.s. p> 0.05, *p<0.05, **p<0.01, ****p<0.0001.

**Extended Data Fig.7, related to Fig. 7.**
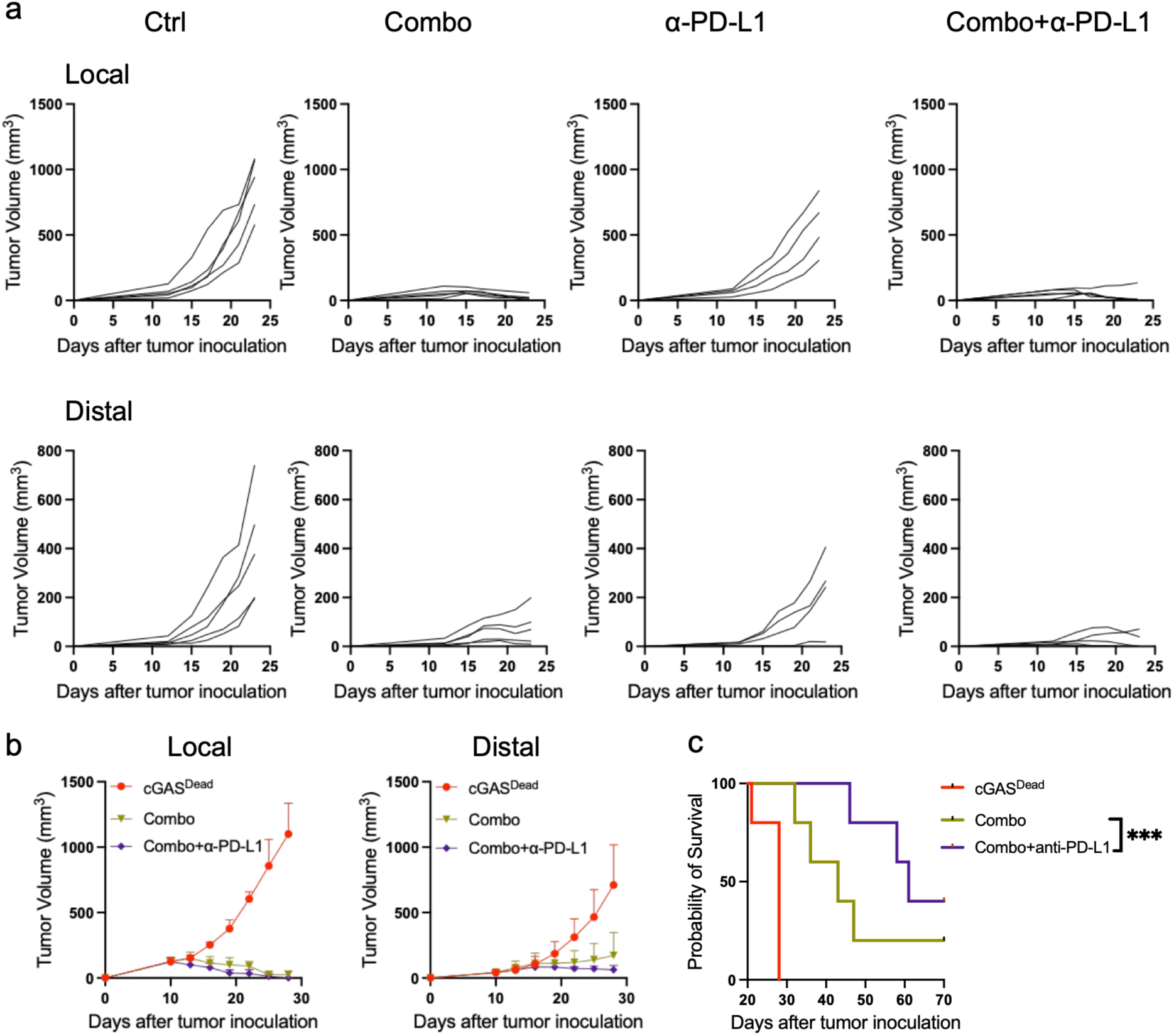
(a) Individual tumor growth curves corresponding to Fig. 7c are shown. (b, c) CT26 tumor bearing mice were treated with Combo LNPs, or additional combination with anti-PD-L1. Tumor growth (b) and overall survival (c) was measured. ***p<0.001.

