## Supplementary Table for "Combining an autoactivated cGAS mutant with membrane-tethered IL-12 via mRNA lipid nanoparticles promotes systemic tumor control"

### Key resource details

| REAGENT or RESOURCE | SOURCE | IDENTIFIER |
| --- | --- | --- |
| **Antibodies** |  |  |
| InVivoMAb anti-mouse CD4 (GK1.5) | BioXcell | Cat# BE0003-1 |
| InVivoMAb anti-mouse CD8 (53-5.8) | BioXcell | Cat# BE0223 |
| InVivoMAb anti-mouse NK1.1 (PK136) | BioXcell | Cat# BE0036 |
| InVivoMAb anti-mouse Ly6C (Monts1) | BioXcell | Cat# BE0203 |
| InVivoMAb anti-mouse Ly6G (1A8) | BioXcell | Cat# BE075-1 |
| InVivoMAb anti-mouse IFN-γ (R4-6A2) | BioXcell | Cat# BE0054 |
| InVivoMAb anti-mouse IFNαR (MAR1-5A3) | BioXcell | Cat# BE0241 |
| anti-PD-L1 (atezolizumab) | Genentech |  |
| Anti-mCD45 (Flow cytometry, 104) | BioLegend | Cat# 109839 |
| Anti-mCD3 (Flow cytometry, KT3.1.1) | BioLegend | Cat# 155614 |
| Anti-mCD4(Flow cytometry, GK1.5) | BioLegend | Cat# 100480 |
| Anti-mCD8α (Flow cytometry, KT15) | Invitrogen | Cat# MA5-16759 |
| Anti-mNK1.1 (Flow cytometry, PK136) | BD Biosciences | Cat# 751111 |
| Anti-mPD-L1(Flow cytometry, MIH5) | BD Biosciences | Cat# 745616 |
| Anti-mCD80(Flow cytometry, 16-10A1) | BD Biosciences | Cat# 741091 |
| Anti-mCD40(Flow cytometry, 3/23) | BD Biosciences | Cat# 741749 |
| Anti-mCD86(Flow cytometry, PO3) | BD Biosciences | Cat# 749021 |
| Anti-mF4/80(Flow cytometry, T45-2342) | BD Biosciences | Cat# 565411 |
| Anti-mI-A/I-E(Flow cytometry, M5/114.15.2) | Biolegend | Cat# 107620 |
| Anti-mCD11c (Flow cytometry, N418) | Biolegend | Cat# 117326 |
| Anti-mXCR1 (Flow cytometry, ZET) | Biolegend | Cat# 148233 |
| Anti-mTox (Flow cytometry, NAN448B) | BD Biosciences | Cat# 569897 |
| Anti-mCXCR3 (Flow cytometry, CXCR3-173) | BD Biosciences | Cat# 757289 |
| Anti-mCD69(Flow cytometry, H1.2F3) | BD Biosciences | Cat# 741234 |
| Anti-mTCF-1(Flow cytometry, S33-966) | BD Biosciences | Cat# 570635 |
| Anti-mFoxp3(Flow cytometry, MF-14) | BD Biosciences | Cat# 126408 |
| Anti-mPD-1(Flow cytometry, BB700) | BD Biosciences | Cat# 566514 |
| Anti-mTim3(Flow cytometry, B8.2C12) | Biolegend | Cat# 134021 |
| Anti-mCD39(Flow cytometry, Duha59) | Biolegend | Cat# 143820 |
| Anti-mKi-67(Flow cytometry, 16A8) | Biolegend | Cat# 652426 |
| Anti-mIL-12 (Flow cytometry, C17.8) | eBioscience | Cat# 12-7123-41 |
| Live dead blue | Invitrogen | Cat# L23105 |
| Fixable Viability Dye eFluor™ 780 | Thermo Fisher | Cat# 65-0865-18 |
| Anti-FcγIII/II receptor (clone 2.4G2) | BD Biosciences | Cat# 553141 |
| **Chemicals, Peptides, and Recombinant Proteins** |  |  |
| IL-2 (SumIL2) “superkine” IL-2 mutations and the F42A mutation<sup>44</sup> (F42A, L80F, R81D, L85V, I86V, and I92F mutants ). | In-house<br>(expressed and purified) | The plasmid is a gift from Dr. Yang-Xin Fu |
| Recombinant Mouse IL-15 Protein | R&D Systems | Cat #: 447-ML |
| Recombinant Mouse IL-15R alpha Fc | R&D Systems | Cat#: 551-MR |
| IL-21 | R&D Systems | Cat #: 594-ML |
| IL-12 Fc | In-house<br>(expressed and purified) |  |
| Recombinant mIFN-γ | R&D Systems | Cat #: 485-MI-100/CF |
| Did | Invitrogen | Cat #: D7757 |
| SM102 | MedKoo | Cat# 464358 |
| DSPC | Avanti | Cat# 850365 |
| DMG-PEG2000 | Avanti | Cat# 880151 |
| Cholestoral | Avanti | Cat# 700100 |
| N1-Methylpseudouridine-5'-Triphosphate | Trilink | Cat# N-1081 |
| CleanCap Reagent AG | Trilink | Cat# N-7113 |
| ATP | Trilink | Cat# N-1510 |
| CTP | Trilink | Cat# N-1511 |
| GTP | Trilink | Cat# N-1512 |
| Murine RNase Inhibitor | New England<br>BioLabs | Cat# M0314 |
| Inorganic Pyrophosphatase | New England<br>BioLabs | Cat# M0361 |
| T7 RNA Polymerase | New England<br>BioLabs | Cat# M0251 |
| TURBO DNase | Invitrogen | Cat# AM2238 |
| Collagenase type I | Sigma | Cat# C0130 |
| DNase I | Roche | Cat# 11284932001 |
| **Critical Commercial Assays** |  |  |
| LEGENDplex™ Mouse Anti-Virus<br>Response Panel (13-plex) | Biolegend | Cat# 740621 |
| Mouse IL-12p70 CBA | BD Biosciences | Cat# 558303 |
| 2'3'-cGAMP ELISA Kit | Cayman<br>Chemical | Cat# 501700 |
| True-Nuclear™ Transcription Factor<br>Buffer Set | Biolegend | Cat# 424401 |
| EasySep™ Mouse TIL (CD45) Positive<br>Selection Kit | STEMCELL | Catalog #100-0350 |
| Chromium Single Cell 5' Library<br>Construction Kit, 16 rxns | 10x Genomics | Cat# PN-1000020 |
| Chromium Next GEM Single Cell 5'<br>Library & Gel Bead Kit v1.1, 16 rxns | 10x Genomics | Cat# PN-1000165 |
| Chromium Single Cell VDJ Enrichment<br>Kit, Mouse T Cell, 96 rxns | 10x Genomics | Cat# PN-1000071 |
| Chromium Next GEM Chip G Single Cell<br>Kit, 16 rxns | 10x Genomics | Cat# PN-1000127 |
| Single Index Kit T Set A, 96 rxns | 10x Genomics | Cat# PN-1000213 |
| Single Index Kit N Set A, 96 rxns | 10x Genomics | Cat# PN-1000212 |
| SPRIselect Reagent Kit | Beckman Coulter | Cat# B23318 |
| High Sensitivity D5000 ScreenTape | Agilent | 5067-5592 |
| High Sensitivity D5000 Reagents | Agilent | 5067-5593 |
| **Software and Algorithms** |  |  |
| GraphPad Prism software 10 | GraphPad Software, Inc. | https://graphpad.com/scientific-software/prism/ |
| SpectroFlo | Cytek | https://cytekbio.com/pages/spectro-flo |
| FlowJo | Tree Star Inc. | https://www.flowjo.com/solutions/flowjo |
| Seurat (v4.1.0) | Satija Lab | https://satijalab.org/seurat/ |
| R (v4.1.1) | R | https://cran.r-project.org |
| Cell Ranger (7.1.0) | 10x Genomics | https://www.10xgenomics.com/ |
| DoubletFinder | DoubletFinder | https://github.com/chris-mcginnis-ucsf/DoubletFinder |
| clusterProfiler | clusterProfiler | https://bioconductor.org/packages/clusterProfiler/ |
